# Non-disruptive 3D profiling of combinations of epigenetic marks in single cells

**DOI:** 10.1101/2025.06.13.659535

**Authors:** Yanbo Chen, Jonathan S.D. Radda, Myles H. Alderman, Shengyan Jin, Yubao Cheng, Yuan Zhang, Miao Liu, Andrew Z. Xiao, Siyuan Wang

**Affiliations:** Department of Genetics, Yale University School of Medicine, New Haven, CT 06510, USA; Yale Stem Cell Center, Yale University School of Medicine, New Haven, CT 06520, USA; Present Address: Department of Cell Biology and Yale Stem Cell Center, Yale University School of Medicine, New Haven, CT, USA; M.D.-Ph.D. Program, Yale University, New Haven, CT 06510, USA; Yale Combined Program in the Biological and Biomedical Sciences, Yale University, New Haven, CT 06510, USA; Molecular Cell Biology, Genetics and Development Program, Yale University, New Haven, CT 06510, USA; Department of Cell Biology, Yale University School of Medicine, New Haven, CT 06510, USA; Biochemistry, Quantitative Biology, Biophysics, and Structural Biology Program, Yale University, New Haven, CT 06510, USA; Yale Center for RNA Science and Medicine, Yale University School of Medicine, New Haven, CT 06510, USA; Yale Liver Center, Yale University School of Medicine, New Haven, CT 06510, USA; Yale Cancer Center, Yale University School of Medicine, New Haven, CT 06510, USA

**Author notes:** These authors contributed equally to this work (Yanbo Chen, Jonathan S.D. Radda, Myles H. Alderman).

## Abstract

Recent advancements in single-cell sequencing and spatial omics technologies have enhanced our understanding of diverse cellular identities, compositions, architectures, and functions. However, the single-cell three-dimensional (3D) organization of the epigenome is still not well understood, due to an absence of spatial single-cell methods that allow high-resolution, locus-specific detection of combinations of epigenetic marks while maintaining the 3D organization of the genome. Here, we develop Epigenetic Proximity Hybridization Reaction (Epi-PHR), a non-disruptive image-based single-cell spatial epigenetic profiling technology. Epi-PHR enables locus-specific and high-resolution *in situ* detection of combinations of epigenetic marks at hundreds of single gene targets within the same individual cells, while retaining the 3D organization of the genome. Dual-mark Epi-PHR in hippocampus tissue sections revealed region-specific epigenetic profiles. Phased Epi-PHR combined with chromatin tracing simultaneously detects allele-specific epigenetic states and chromatin conformations of a paternally imprinted *Meg3* gene cluster in single cells, revealing associations between specific epigenetic mark enrichment and chromatin folding features for the distinct alleles from different parental origins. Surprisingly, we found that H3K9me3 abundance is positively associated with a chromatin domain boundary at the *Meg3* locus among maternal copies. We expect Epi-PHR will be broadly applicable in research requiring single-cell spatial epigenetic information, and will help extend our understanding of the combinatorial epigenetic code and its relationship with chromatin organization.

## Main

In eukaryotes, the functional outputs of genomes are intricately regulated by local epigenetic states encoded by a variety of epigenetic marks and three-dimensional (3D) chromatin folding^1–5^. These regulatory mechanisms are crucial in controlling many genomic and cellular functions and processes, e.g. transcription, DNA replication timing, and DNA damage response^1–9^. The epigenome and the 3D genome are also closely interlinked: For example, DNA methylation and histone modifications are known to alter local chromatin structure, thereby influencing whether specific genomic regions are accessible for transcription and other cellular processes^10,11^.

Multiple recently discovered 3D genome architectural units, such as A-B compartments, lamina-associated domains (LADs), and nucleolus-associated chromatin domains (NADs) are enriched with active or inactive epigenetic marks^4,12–16^. Topologically associating domain (TAD, also known as contact domain^17–21)^ boundaries, often bound by insulator protein CTCF, sometimes demarcate epigenetic domains^17,19^. Furthermore, chromatin folding has been proposed to help maintain epigenetic memory^22,23^.

Traditionally, distinct sequencing-based methods are employed to separately profile genome-wide epigenetic states and chromatin folding in ensembles of cells, which only provide population-averaged information^12,24–28^. More recently, significant advancements have been made in sequencing-based techniques to enable the characterization of epigenetic modifications and chromatin folding at the single-cell level^29–39^. Recent spatial epigenomic sequencing methods also enabled epigenetic profiling with near-single-cell spatial resolution^40,41^. However, sequencing-based single-cell epigenetic mark and 3D genome co-profiling methods are limited to profiling DNA methylation together with the 3D genome^37,38^. In addition, sequencing-based single-cell or spatial histone modification profiling techniques often suffer from low sequencing read count per cell or per spatial spot, limiting gene-locus-specific analyses at the single-cell level.

In contrast to sequencing-based methods, imaging-based methods are an attractive alternative as they by default preserve the spatial information of the biological sample. Coimmunofluorescence of epigenetic marks is compatible with highly multiplexed DNA fluorescence *in situ* hybridization (FISH) targeting numerous genomic loci, but often suffers from high background and low resolution of the antibody staining, again limiting gene-locus-specific readout of epigenetic states^42,43^. A new image-based epigenetic profiling method, termed epigenomic multiplexed error-robust FISH (epigenomic MERFISH), is the only *in situ* method that allows high genomic resolution (2 kb) and locus-specific readout of epigenetic states of numerous genomic loci in single cells^44^. However, epigenomic MERFISH tends to disrupt the 3D genome, as demonstrated in this work, has poor performance when profiling heterochromatic mark, and is currently limited to profiling one epigenetic mark at a time.

In this work, we developed the first non-disruptive, high resolution, *in situ* epigenetic profiling method. Unlike epigenomic MERFISH, our method retains 3D genome information, allows for high-performance heterochromatic mark profiling, and enables mapping of combinations of epigenetic marks at numerous genomic loci. We demonstrated the ability of our method to profile epigenetic marks at genomic regions as small as 2 kb, allowing the method to target many single genes and short genomic elements. We also demonstrated the multiplexing capability of our method by combining it with a barcoding strategy to profile the epigenetic states and nuclear distributions of hundreds of genes in the same single cells, both for single epigenetic marks and combinations of marks, in cell cultures and in mouse hippocampus tissue sections. Finally, we combined our method with allelic DNA FISH and chromatin tracing to simultaneously detect parental allele-specific epigenetic state, chromatin folding conformation, and nuclear distribution of an imprinted cluster in the same single cells of a hybrid mouse trophoblast stem cell (TSC) model, revealing unexpected information that cannot be acquired with existing allele-specific 3D genome and epigenome sequencing approaches.

## Results

### Epi-PHR can robustly detect epigenetic marks in single cells

To detect specific histone modifications at a target genomic locus *in situ*, here we developed Epigenetic Proximity Hybridization Reaction (Epi-PHR) (Fig. 1a), in part inspired by proximity-dependent hybridization chain reaction (proxHCR)^45,46^ that detects protein-protein interactions. In Epi-PHR, fixed and permeabilized cell samples were incubated first with a primary antibody specific to the epigenetic mark of interest, followed by a biotinylated secondary antibody. After post-fixation, the samples were embedded in polyacrylamide gels and then incubated with genomic-locus-specific FISH probes. Then the core Epi-PHR mechanism was initiated by the spatial proximity of two DNA hairpin oligonucleotides, PH1 and PH2. PH1 was hybridized to the DNA FISH probes through a docking adapter, and biotinylated PH2 was linked to the biotinylated antibodies through streptavidin. The proximity required for Epi-PHR was achieved when the antibodies bound to the epigenetic marks were spatially adjacent to the FISH probes bound to the DNA locus. An activator oligonucleotide was then introduced to the system, which hybridized with PH1 and caused its hairpin to open. The opened PH1 then invaded and opened a nearby PH2 hairpin. Subsequently, a fluorescent dye-labeled oligonucleotide probe, H1, was hybridized to the opened end of PH2. Instead of using uncontrolled hybridization chain reaction signal amplification^45,46^, we sequentially incubated a second dye-labeled oligonucleotide probe, H2, to achieve a controlled two-fold signal amplification. This approach generates a fluorescent readout (Epi-PHR signal) that indicates the presence of an epigenetic mark of interest at a specific genomic locus (Fig. 1a).

**Fig. 1.**
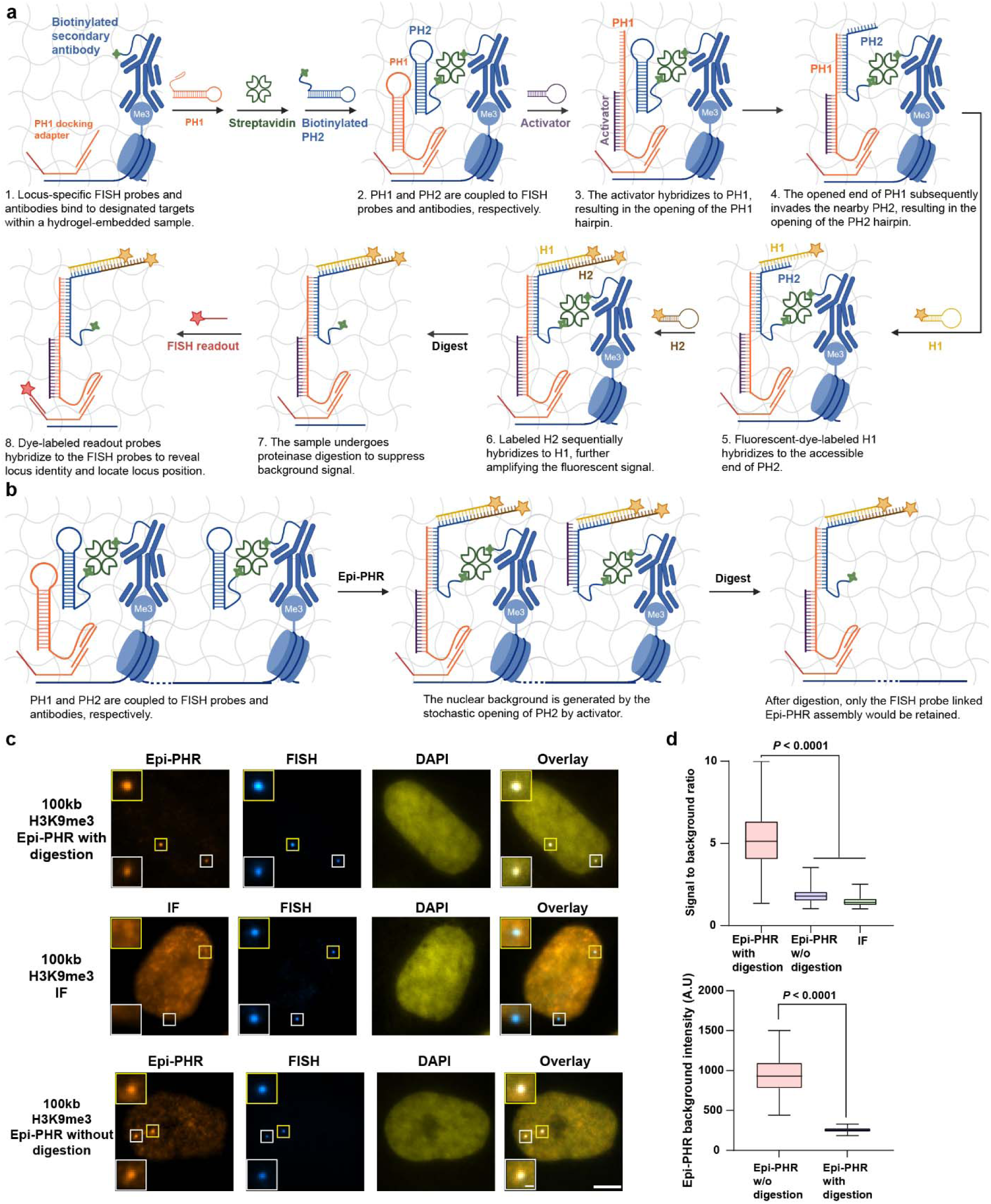
| Locus specific epigenetic mark detection by Epi-PHR. **a**, Detailed schematic illustration of Epi-PHR for locus specific epigenetic mark detection. **b**, Schematic illustrations of the source of Epi-PHR background signal, and how proteinase digestion addresses high background problem. **c**, Representative H3K9me3 IF, FISH, Epi-PHR, and DAPI images using a 100-kb FISH probe library. IF signals were generated by dye-labeled streptavidin. The corner panels show magnified images that use FISH signals as center positions. Scale bar for the original images: 5 µm; scale bar for the magnified images: 500 nm. **d**, Comparing the signal-to-background ratios of Epi-PHR and IF, and the effect of proteinase K digestion on Epi-PHR background. The signal-to-background ratios for Epi-PHR and IF were measured at FISH positions. For each cell, the median fluorescent intensity in the Epi-PHR color channel within the DAPI-defined nuclear region was used as the Epi-PHR background measurement. In box plots, central lines represent median values, boxes show interquartile ranges (IQRs), and whiskers indicate non-outlier minimum and maximum values. Outliers were defined as values further than 1.5 times the IQR away from the box. Statistical significance was determined by unpaired two-tailed Mann-Whitney test. Sample sizes are as follows: Epi-PHR signal-to-background ratio with digestion (n = 365), Epi-PHR signal-to-background ratio without digestion (n = 343), IF signal-to-background ratio (n = 292), Epi-PHR background intensity with digestion (n = 219), Epi-PHR background intensity without digestion (n = 198).

During the development of Epi-PHR, we observed that high concentrations of the activator could stochastically open PH2, leading to the generation of an Epi-PHR background signal that closely resembled the pattern seen in immunofluorescence (IF) staining (Fig. 1b). This unintended activation presented challenges in distinguishing the locus-specific signals of interest from background noise, especially when targeting small genomic loci. To address this issue, we implemented a protocol wherein samples were first embedded into polyacrylamide gels following antibody incubation. After completing the Epi-PHR procedure, we subjected the samples to proteinase K digestion^47^. This procedure digested antibodies and removed their associated PH2 oligos, unless the PH2 oligos were coupled with PH1 oligos. This strategy ensured that only the PH1-linked Epi-PHR assemblies, which were physically associated with the genomic-locus-specific FISH probes, were retained after the digestion (Figs. 1a and 1b), and the strategy effectively removed the background signal associated with stochastic PH2 activation (Figs. 1c and 1d). Finally, to determine the 3D position of the target DNA locus, we hybridized a dye-labeled readout probe to the locus-specific FISH probes and visualized the DNA locus (FISH signal) in a second color channel (Fig. 1a).

As a proof-of-principle demonstration of Epi-PHR, we targeted a 100-kb region on human chromosome 1 (chr1: 4742385 – 4842385, hg19) in cultured IMR90 cells. In these cells, the target locus is known to be enriched with H3K9me3, based on published chromatin immunoprecipitation sequencing (ChIP-seq) data^48^. Following an Epi-PHR procedure with H3K9me3 antibodies, we observed strong Epi-PHR signals colocalizing with FISH signals (Fig. 1c, and Extended Data Fig. 1a, first row), which indicated the detection of H3K9me3 at the targeted locus. To compare Epi-PHR with IF staining, we used dye-labeled streptavidin to visualize IF signals (Fig. 1c, second row). The IF signals exhibited a whole nuclear staining pattern, with IF signals at FISH positions only slightly above background levels (Fig. 1d, top panel). In contrast, Epi-PHR generated distinct foci-like signals even without proteinase digestion (Fig. 1c, third row), consistent with the locus-specific signal generation of the Epi-PHR system. After digestion, the nuclear background of Epi-PHR was significantly reduced (Fig. 1c, first row, and 1D, bottom panel), further enhancing the Epi-PHR signal-to-background ratio (Fig. 1d, top panel).

To further validate the Epi-PHR system, we conducted systematic negative control experiments by omitting individual key Epi-PHR components to verify the necessity of each component (Extended Data Fig. 1a). In these experiments, we observed robust H3K9me3 Epi-PHR signals only when all components were present. Conversely, the omission of any single Epi-PHR component, including the activator oligonucleotide, H1, PH1, PH2, secondary antibody, and streptavidin, completely abolished H3K9me3 Epi-PHR signals, demonstrating the dependency of the system on each individual component for robust Epi-PHR signal generation (Extended Data Fig. 1b). These results demonstrate that the Epi-PHR system operates as designed for detecting epigenetic modifications.

### Epi-PHR allows specific and high-genomic-resolution detection of epigenetic marks in single cells

To determine the sensitivity of the Epi-PHR system at different genomic resolutions, we systematically decreased the size of the target genomic region and reduced the number of FISH probes used to tile the region (Fig. 2a), scaling from 100 kb (2219 different FISH probes) down to 2 kb (52 different FISH probes). Target coordinates were carefully selected to ensure uniform scaling of FISH probe count and H3K9me3 ChIP-seq^48^ peak coverage (Fig. 2b). Across all tested target lengths, we observed strong Epi-PHR signals that colocalized with FISH foci, whereas negative control samples without primary antibodies showed no visible Epi-PHR signal (Fig. 2c). At all tested resolutions, we achieved a high FISH detection efficiency of ∼2 foci per cell in this diploid cell line (Fig. 2d). We observed a gradual decrease in Epi-PHR signal-to-background ratio as the targeted regions were progressively reduced in size (Fig. 2e). Despite the varying signal-to-background ratios, the Epi-PHR signal detection efficiency was high at all resolutions, ranging from 99% at 100 kb resolution to 59% at 2 kb resolution (1% false positive for the control without primary antibody) (Fig. 2e). This observation suggests that Epi-PHR signal intensity is dependent on the number of FISH probes and the quantity of the histone mark present within the targeted region. Furthermore, the clear Epi-PHR signal detected at 2 kb target length indicates the potential for detecting epigenetic marks associated with single genes or genomic elements. Overall, these results demonstrate the capability of Epi-PHR to detect epigenetic marks across different genomic length scales.

**Fig. 2.**
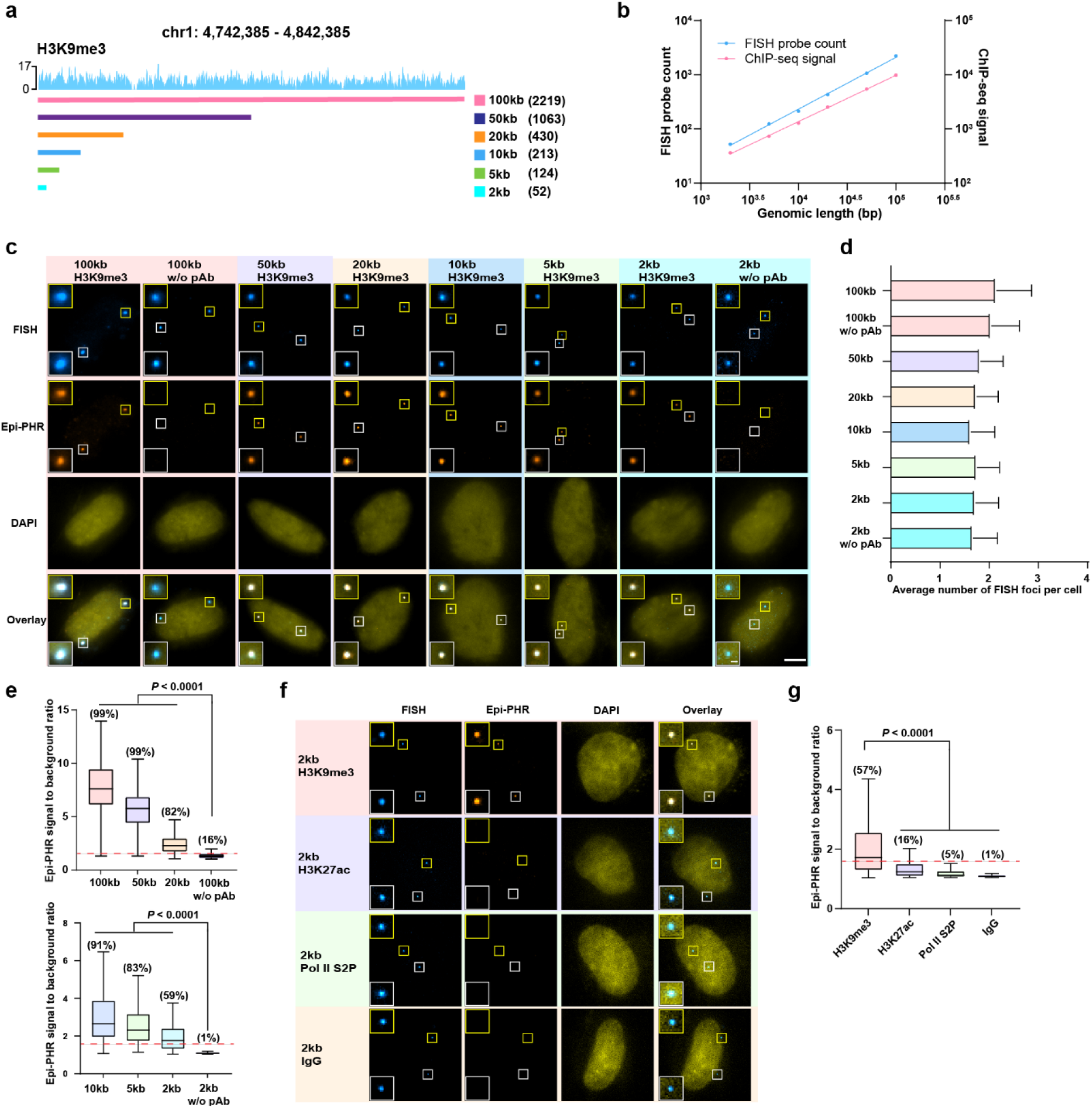
| Highly sensitive and specific detection of epigenetic marks by Epi-PHR. **a**, Schematic illustration of FISH probe coverage for Epi-PHR sensitive test alongside with the H3K9me3 ChIP-seq data from targeted region^48^. The number in the parentheses indicates the total number of different FISH probes for each tested region. **b**, FISH probe count (blue points) and ChIP-seq signal^48^ (red points) versus genomic length for targeted regions. Lines represent linear regression results. **c**, Representative H3K9me3 FISH, Epi-PHR, and DAPI images using different genomic resolutions of FISH probe libraries. Control samples omitted primary antibody and used 100-kb and 2-kb probe sets. All color channels were used to generate overlay images. The corner panels show magnified images that use FISH signals as center positions. Scale bar for the original images: 5 µm; scale bar for the magnified images: 500 nm. **d**, Number of detected FISH spots per cell. Bars represent mean values and whiskers extend to standard deviations. **e**, Signal-to-background ratios of Epi-PHR measured at FISH positions using different sets of FISH probes. Detection efficiencies are in parentheses. The dashed line indicates the defined detection efficiency threshold (Methods). In box plots, central lines represent median values, boxes show interquartile ranges (IQRs), and whiskers indicate non-outlier minimum and maximum values. Outliers were defined as values further than 1.5 times the IQR away from the box. Statistical significance was determined by unpaired two-tailed Mann-Whitney test. Sample sizes are as follows: 100-kb probe set (n = 217), 50-kb probe set (n = 173), 20-kb probe set (n = 162), 100-kb without primary antibody (n = 241), 10-kb probe set (n = 127), 5-kb probe set (n = 177), 2-kb probe set (n = 181), 2-kb without primary antibody (n = 137). **f**, Representative FISH, Epi-PHR, and DAPI images using H3K9me3, H3K27ac, Pol II S2P, or rabbit IgG control antibody, using the 2-kb FISH probe set. All color channels were used to generate overlay images. The corner panels show magnified images that use FISH signals as center positions. Scale bar for the original images: 5 µm; scale bar for the magnified images: 500 nm. **g**, Signal-to-background ratios of Epi-PHR measured at FISH positions using H3K9me3, H3K27ac, Pol II S2P, or rabbit IgG control antibody. Detection efficiencies are in parentheses. The dashed line indicates the detection efficiency threshold. In box plots, central lines represent median values, boxes show interquartile ranges (IQRs), and whiskers indicate non-outlier minimum and maximum values. Outliers were defined as values further than 1.5 times the IQR away from the box. Statistical significance was determined by an unpaired two-tailed Mann-Whitney test. Sample sizes are as follows: H3K9me3 antibody (n = 274), H3K27ac antibody (n = 263), Pol II S2P antibody (n = 249), rabbit IgG control antibody (n = 214).

To further validate the specificity of the Epi-PHR system, we targeted active marks H3K27ac and RNA polymerase II (Pol II) S2P within the same 2 kb region described above, as well as with a nonspecific IgG antibody as the primary antibody. Because this region was situated within a large stretch of H3K9me3 enriched heterochromatin^48^, we hypothesized only H3K9me3 Epi-PHR would produce strong signals. Indeed, strong Epi-PHR signals were only observed when targeting H3K9me3, and H3K9me3 Epi-PHR detection efficiency was significantly higher than those of H3K27ac, Pol II S2P, and IgG control (Figs. 2f and 2g). The remaining positive detections of H3K27ac and Pol II S2P above the IgG control may be in part attributed to limited specificity of the antibodies targeting these marks. Overall, the results here demonstrate the high specificity of the Epi-PHR method.

### Epi-PHR can profile epigenetic marks at hundreds of genomic loci in single cells

We next aimed to increase the multiplexing capacity and genomic throughput of our system first by increasing the number of target genomic loci. Towards this end, we combined Epi-PHR with DNA multiplexed error robust FISH (DNA MERFISH)^49^, to target and distinguish numerous genomic loci in the same single cells (Fig. 3a). We designed a library of primary FISH probes targeting 200 non-repetitive gene loci, with each probe consisting of a primary targeting sequence, a PH1 docking sequence, and two readout-probe-binding regions that allow visualization via dye-labeled readout oligonucleotide probes. High multiplexing was achieved by assigning different combinations of readout-probe-binding regions to each group of primary probes targeting a specific genomic locus of interest. The presence or absence of different readout-probe-binding regions on the primary probes targeting each locus formed a unique binary barcode that encoded the target locus identity, with the presence or absence of the regions corresponding to binary digit 1 or 0, respectively. Because all FISH probes contained the same PH1 docking sequence, Epi-PHR signals for all target loci were generated simultaneously. After imaging the Epi-PHR signals, dye-labeled readout probes were sequentially introduced to the sample to visualize and read out the barcodes of target loci based on the presence or absence of FISH signals across the sequential imaging rounds. The genomic identities of the target loci were then decoded from the imaged barcodes. Between sequential rounds of readout probe hybridization and imaging, the fluorescent dyes on the already imaged readout probes were removed by chemical cleavage before the next round of readout hybridization. In our design, we assigned each target locus a unique 36-bit, Hamming weight 2 (HW2) barcode (i.e. each code contains 36 binary digits and two “1” digits), and read out these barcodes using 18 sequential rounds of two-color imaging (Fig. 3a). The Epi-PHR images were aligned with the multiplexed FISH images to measure the Epi-PHR signals at the decoded FISH foci (Fig. 3b).

**Fig. 3.**
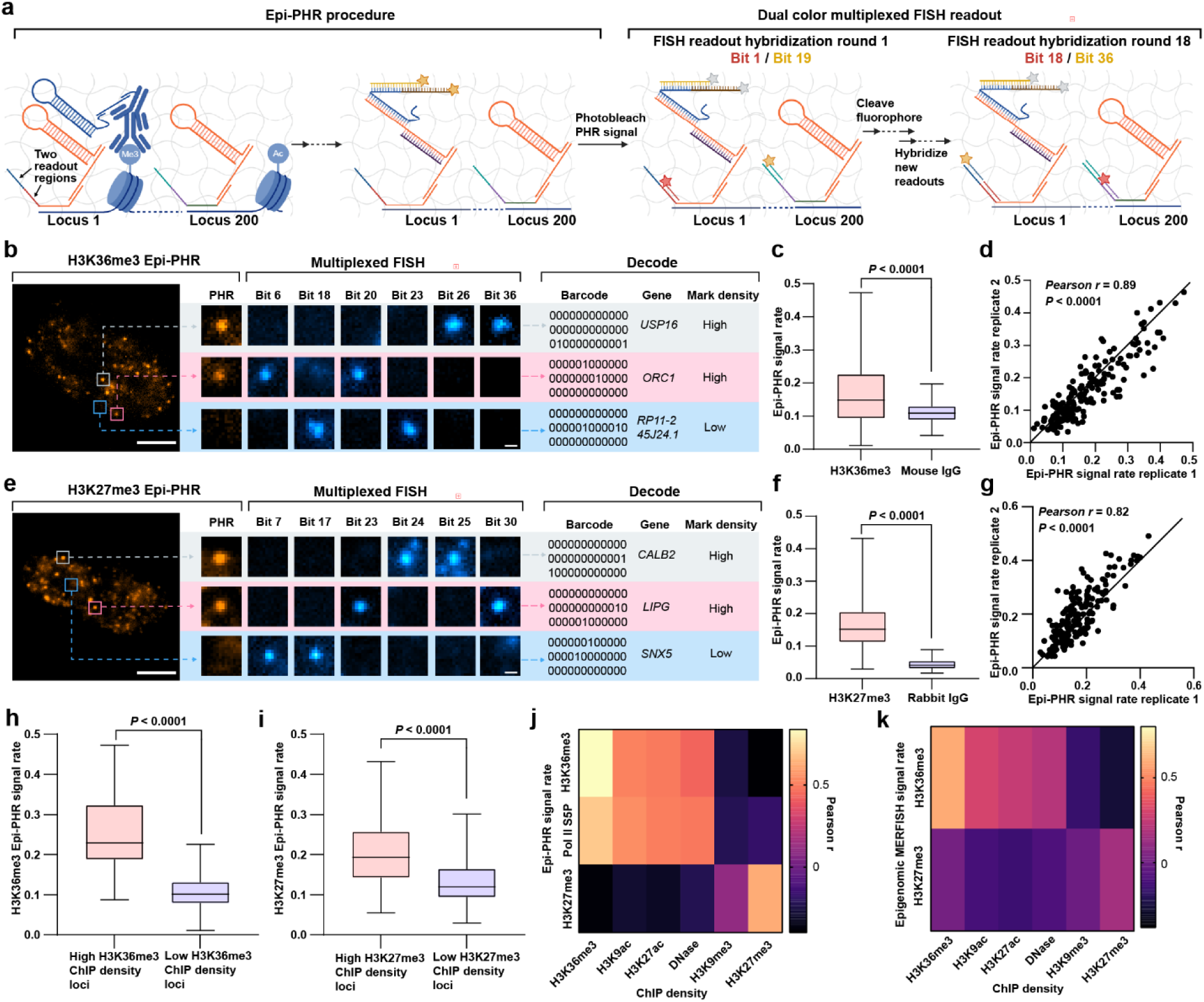
| Multi-loci epigenetic mark profiling by Epi-PHR. **a**, Schematic illustration of multiplexed Epi-PHR design. PH1 and PH2 hybridize to different adapters that recognize the FISH probe and oligo-conjugated secondary antibody, respectively. Following the Epi-PHR procedure, signals from all targeted loci were recorded. After photobleaching the Epi-PHR signals, 18 rounds of two-color imaging were conducted to capture all barcoded loci. **b**, Representative 200-gene H3K36me3 Epi-PHR and multiplexed FISH images. Magnified images display three decoded loci with their associated Epi-PHR signals and multiplexed FISH signals in selected rounds of readout hybridization. Scale bar for the original images: 5 µm; scale bar for the magnified images: 500 nm. **c**, Epi-PHR signal rates using H3K36me3 antibody or mouse IgG control antibody. The H3K36me3 Epi-PHR signal threshold was used for Epi-PHR signal rates calculation for both samples. Only genes with more than 60 measurements are included in this analysis. In box plots, central lines represent median values, boxes show interquartile ranges, and whiskers indicate minimum and maximum values. Statistical significance was determined by a Wilcoxon paired signed rank test. Sample sizes are as follows: H3K36me3 antibody (n = 183 genes), mouse IgG antibody (n = 183 genes). **d**, Epi-PHR signal rate correlation between two biological replicates of H3K36me3 Epi-PHR experiments. The Pearson correlation coefficient and statistical significance determined by two-tailed test are shown. **e**, Representative 200-gene H3K27me3 Epi-PHR and multiplexed FISH images. Magnified images display three decoded loci with their associated Epi-PHR signals and multiplexed FISH signals in selected rounds of readout hybridization. Scale bar for the original images: 5 µm; scale bar for the magnified images: 500 nm. **f**, Epi-PHR signal rates using H3K27me3 antibody or rabbit IgG control antibody. The H3K27me3 Epi-PHR signal threshold was used for Epi-PHR signal rates calculation for both samples. Only genes with more than 60 measurements are included in this analysis. In box plots, central lines represent median values, boxes show interquartile ranges, and whiskers indicate minimum and maximum values. Statistical significance was determined by a Wilcoxon paired signed rank test. Sample sizes are as follows: H3K27me3 antibody (n = 187 genes), rabbit IgG antibody (n = 187 genes). **g**, Epi-PHR signal rate correlation between two biological replicates of H3K27me3 Epi-PHR experiments. The Pearson correlation coefficient and statistical significance determined by two-tailed test are shown. **h-i**, Box plots quantifying the Epi-PHR signal rates by categorizing targeted loci into high or low ChIP-seq peak density groups. For each targeted locus, ChIP-seq^48^ density is calculated by summing peak heights over the targeted region and normalizing to the genomic length of the targeted region. The high ChIP-seq peak density group includes loci with a ChIP-seq density greater than zero. Only loci with more than 60 Epi-PHR measurements are included in this analysis. In box plots, central lines represent median values, boxes show interquartile ranges, and whiskers indicate minimum and maximum values. Statistical significance was determined by an unpaired two-tailed Mann-Whitney test. Sample sizes are as follows: high H3K36me3 ChIP density loci (n = 82 genes), low H3K36me3 ChIP density loci (n = 101 genes), high H3K27me3 ChIP density loci (n = 102 genes), low H3K27me3 ChIP density loci (n = 85 genes). **j**, Pearson correlation matrix comparing H3K36me3, Pol II S5P, and H3K27me3 Epi-PHR signal rates with H3K36me3, H3K9ac, H3K27ac, DNase, H3K9me3, and H3K27me3 ChIP-seq densities^48^. See Extended Data Fig. 2c for correlation plots. **k**, Pearson correlation matrix comparing H3K36me3 and H3K27me3 epigenomic MERFISH signal rates with H3K36me3, H3K9ac, H3K27ac, DNase, H3K9me3, and H3K27me3 ChIP-seq densities^48^. See Extended Data Fig. 2g for correlation plots.

We first profiled H3K36me3 at the 200 selected gene loci in cultured IMR90 cells. H3K36me3 is a canonical mark associated with active transcription, deposited through elongating RNA Pol II complex to maintain transcriptional fidelity and prevent cryptic transcription initiation within gene bodies^50–52^. From the raw images, we observed clear foci-like Epi-PHR signals in individual nuclei, and multiplexed FISH showed distinct foci across different rounds of hybridization, allowing decoding of the gene identities of the FISH foci (Fig. 3b). To demonstrate that the H3K36me3 antibody specifically generated the Epi-PHR signals, we also profiled these 200 gene loci using a mouse control IgG antibody. Comparing the Epi-PHR signal rates, defined as the probability of observing high Epi-PHR signal over background at each target locus (Methods), we observed significantly lower Epi-PHR signal rates when using the control IgG antibody (Fig. 3c). Moreover, the H3K36me3 Epi-PHR signal rates showed no correlation with the control IgG Epi-PHR signal rates (Extended Data Fig. 2a), confirming that the observed H3K36me3 Epi-PHR signals were not due to antibody nonspecific binding.

Furthermore, the H3K36me3 Epi-PHR results were highly reproducible, with a Pearson correlation coefficient of 0.89 between biological replicates (Fig. 3d). These findings validate the specificity and reproducibility of the Epi-PHR system in detecting epigenetic marks at numerous genomic loci.

Using the same library, we next profiled the repressive mark H3K27me3 associated with transcriptional silencing^53^ at the same 200 gene loci. Epi-PHR signals co-localized with multiplexed FISH signals were observed within the nuclei (Fig. 3e). The H3K27me3 Epi-PHR signal rates were significantly higher than those obtained with the rabbit control IgG (Fig. 3f). Furthermore, there was no correlation between the Epi-PHR signal rates of H3K27me3 and the control IgG, supporting the specificity of the observed H3K27me3 Epi-PHR signals (Extended Data Fig. 2b). The reproducibility of the H3K27me3 Epi-PHR results was also high, with a Pearson correlation coefficient of 0.82 (Fig. 3g). These results further demonstrate the specificity and reliability of the Epi-PHR system.

As a further test, we cross-validated the Epi-PHR results with publicly available ChIP-seq data^48^. We categorized the target loci into a high ChIP-seq peak density group and a low ChIP-seq peak density group (Methods). For both H3K36me3 and H3K27me3 marks, we observed higher Epi-PHR signal rates among the corresponding high ChIP-seq peak density loci, indicating a general agreement between the two approaches (Figs. 3h and 3i). Furthermore, we correlated the Epi-PHR signal rates with various ChIP-seq datasets (Fig. 3j). Notably, the Epi-PHR signal rates showed strong correlations with ChIP-seq data of the same epigenetic mark, with Pearson correlation coefficients of 0.84 for H3K36me3 and 0.62 for H3K27me3 (Fig. 3j and Extended Data Fig. 2c). Additionally, we found that H3K36me3 Epi-PHR signal rate also correlated with ChIP-seq peak densities of other active marks, such as H3K27ac (Fig. 3j and Extended Data Fig. 2c), consistent with previous reports that H3K36me3 and H3K27ac are marks for actively transcribed genes^1,48^. Conversely, there was low negative correlation between H3K36me3 Epi-PHR signal rates and ChIP-seq densities of repressive marks like H3K9me3 and H3K27me3 (Fig. 3j and Extended Data Fig. 2c). Interestingly, we did not observe strong correlations between the H3K27me3 Epi-PHR signal rate and the ChIP-seq densities of other marks, including the repressive mark H3K9me3. This result aligns with previous studies indicating that facultative and constitutive heterochromatin, marked by H3K27me3 and H3K9me3, respectively, play distinct roles in maintaining genomic integrity and regulating gene expression^54,55^.

Given that H3K36me3 deposition depends on actively transcribing RNA Pol II complex, we hypothesized that the Epi-PHR profile of RNA Pol II S5P, a modification indicating active Pol II, would be similar to that of H3K36me3. To test this hypothesis, we also profiled RNA Pol II S5P (Extended Data Fig. 2d) and found that the RNA Pol II S5P Epi-PHR signal rate was correlated with the H3K36me3 ChIP-seq^48^ peak density, with a Pearson correlation coefficient of 0.70 (Fig. 3j and Extended Data Fig. 2c). Additionally, RNA Pol II S5P Epi-PHR signal rate was highly correlated with H3K36me3 Epi-PHR signal rate (Extended Data Fig. 2e), consistent with the co-occupancy of these two marks^56^. These results further validate the Epi-PHR system’s robustness and demonstrate its versatility in profiling various epigenetic components beyond histone modifications, including the transcription machinery.

With the preserved nuclear organization in Epi-PHR (see a later section for detailed analysis), we were able to explore the relationship between the nuclear distribution of genomic loci and their epigenetic states. To achieve this, we extracted the spatial locations of loci with high Epi-PHR signals and measured the distance between these loci and the nuclear periphery. We observed that H3K27me3 enriched loci were in closer proximity to the nuclear periphery compared to H3K36me3 enriched loci (Extended Data Fig. 2f). This finding aligns with previous observations that H3K27me3 enriched loci have increased association with the nuclear periphery^57^. These results demonstrate that Epi-PHR allows sub-nuclear spatial mapping of the epigenetic states of numerous genomic loci.

To further compare Epi-PHR with state-of-the-art image-based spatial epigenetic profiling technology, we performed epigenomic MERFISH^44^ experiments targeting H3K36me3 and H3K27me3 at the same 200 gene loci. The epigenomic MERFISH signal rate of the active mark H3K36me3 exhibited a good (albeit lower than Epi-PHR) correlation with ChIP-seq data, with a Pearson correlation coefficient of 0.59 (Fig. 3k and Extended Data Fig. 2g). In contrast, the repressive mark H3K27me3 epigenomic MERFISH data had a substantially lower correlation with ChIP-seq data, with a Pearson correlation coefficient of 0.15 (Fig. 3k and Extended Data Fig. 2g), potentially attributable to a likely low tagmentation efficiency in highly compacted heterochromatin regions with the fixation condition needed for epigenomic MERFISH. These results highlight the advantage of Epi-PHR in its capability to target repressive epigenetic marks.

### Epi-PHR can simultaneously profile more than one epigenetic mark at hundreds of genomic loci in single cells

To further extend the multiplexing capability of Epi-PHR, we developed a strategy to simultaneously profile pairs of epigenetic marks in the same single cells (Fig. 4a). Using previously published methods^45,46^, we designed a second set of Epi-PHR hairpins (PH1, PH2, activator, H1 and H2), orthogonal to the first set, that could be used to target a second epigenetic mark. The second set of H1 and H2 hairpins was labeled with a dye different from the first set of hairpins to enable parallel visualization in a second color channel. To prevent cross-talk between the two hairpin sets and ensure their uniform incubation conditions, the structure and melting temperature of the second set of hairpins were closely matched to those of the first set. To generate mark-specific Epi-PHR signals, we first selected pairs of primary antibodies from distinct species targeting different epigenetic marks. We then linked the two PH2 hairpins to different secondary antibodies that would selectively bind to the primary antibodies. To enable opening of either PH2 hairpin when in proximity, locus-specific FISH probes docked both PH1 hairpins through an adapter. Upon triggering Epi-PHR with the two activator oligos, the sets of distinct H1 and H2 oligos were introduced to generate Epi-PHR signals for both of the targeted epigenetic marks in separate color channels (Fig. 4a). As in the one-mark profiling experiments, loci identities and their 3D positions were read out by multiplexed FISH.

**Fig. 4.**
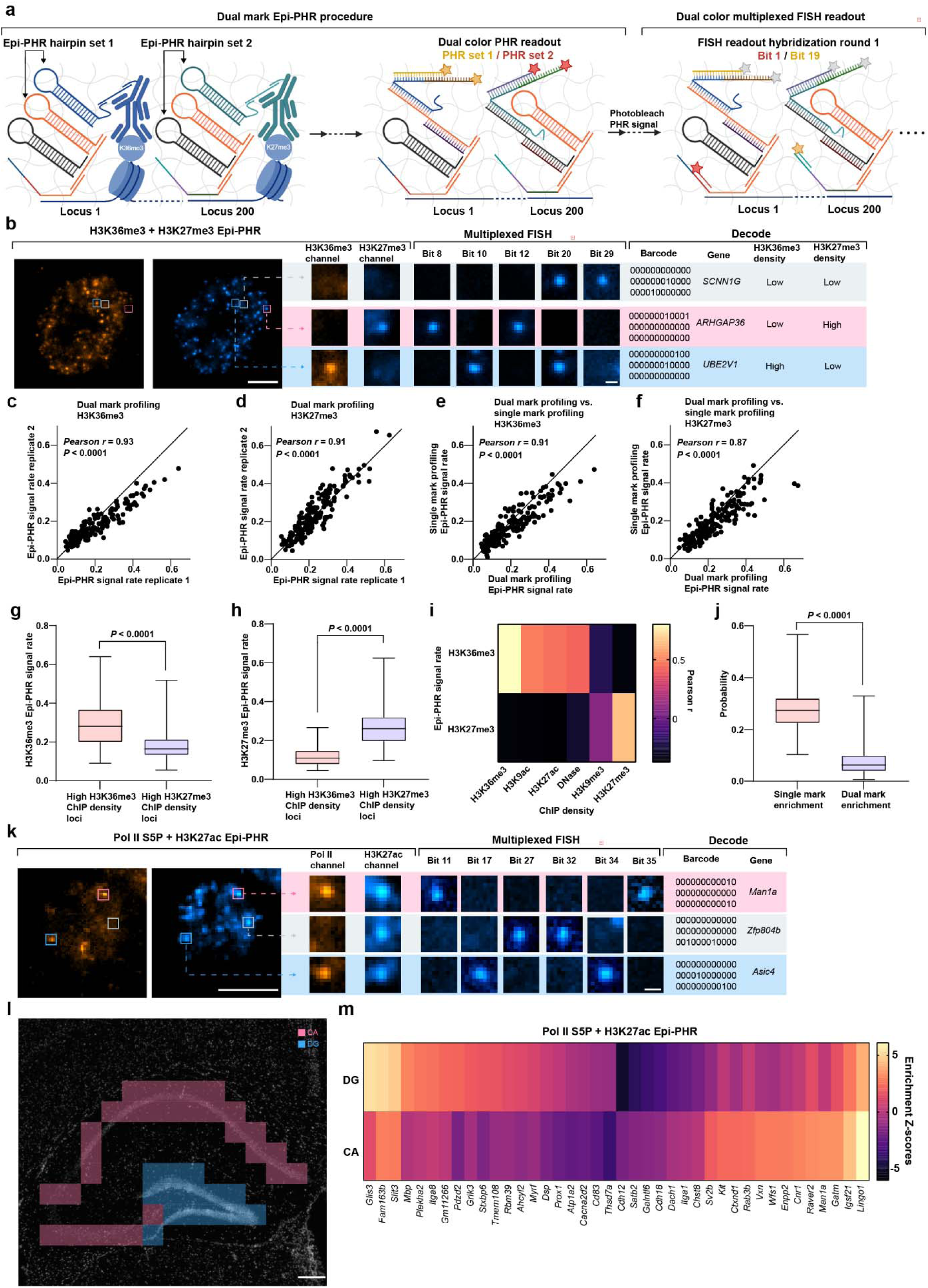
| Dual epigenetic mark profiling by Epi-PHR. **a**, Schematic illustration of dual-mark Epi-PHR design. PH1 oligos from both sets hybridize to FISH probes via an adapter. PH2 oligos hybridize to different adapters that recognize the oligo-conjugated mouse secondary antibody (H3K36me3) and rabbit secondary antibody (H3K27me3), respectively. Following the Epi-PHR procedure, H3K27me3 and H3K36me3 Epi-PHR signals from all targeted loci were recorded in different color channels. After photobleaching the Epi-PHR signals, 18 rounds of two-color imaging were conducted to capture all barcoded loci. **b**, Representative 200-gene H3K36me3-H3K27me3 dual-mark Epi-PHR and multiplexed FISH images. Magnified images display three decoded loci with their associated Epi-PHR signals in different Epi-PHR channels and multiplexed FISH signals in selected rounds of readout hybridization. Scale bar for the original images: 5 µm; scale bar for the magnified images: 500 nm. **c-d**, Epi-PHR signal rate correlation between two biological replicates of H3K36me3-H3K27me3 dual-mark Epi-PHR experiments. Pearson correlation coefficients and statistical significance determined by two-tailed test are shown. **e-f**, Epi-PHR signal rate correlation between single-mark profiling and dual-mark profiling. Pearson correlation coefficients and statistical significance determined by two-tailed test are shown. **g-h**, Box plots quantifying the Epi-PHR signal rates by categorizing targeted loci into high or low ChIP-seq peak density group. The categorization criteria are described in Figs. 3h and 3i, and only loci with more than 60 Epi-PHR measurements are included in this analysis. In box plots, central lines represent median values, boxes show interquartile ranges, and whiskers indicate minimum and maximum values. Statistical significance was determined by an unpaired two-tailed Mann-Whitney test. Sample sizes are as follows: high H3K36me3 ChIP density loci (n = 85 genes), low H3K36me3 ChIP density loci (n = 101 genes), high H3K27me3 ChIP density loci (n = 101 genes), low H3K27me3 ChIP density loci (n = 85 genes). **i**, Pearson correlation matrix comparing H3K36me3-H3K27me3 dual-mark Epi-PHR profiling signal rates with H3K36me3, H3K9ac, H3K27ac, DNase, H3K9me3, H3K27me3 ChIP-seq densities^48^. See Extended Data Fig. 4a for correlation plots. **j**, Probability of single mark enrichment and dual mark enrichment for each targeted locus. The probability of dual mark enrichment is calculated by dividing the number of foci with Epi-PHR signals greater than both H3K36me3 and H3K27me3 thresholds by the total number of decoded foci of the targeted locus. The probability of single mark enrichment is calculated by dividing the number of foci with Epi-PHR signals greater than one threshold but smaller than the other by the total number of decoded foci of the targeted locus. In box plots, central lines represent median values, boxes show interquartile ranges, and whiskers indicate minimum and maximum values. Statistical significance was determined by a Wilcoxon paired signed rank test. Sample sizes are as follows: single mark enrichment (n = 186 genes), dual mark enrichment (n = 186 genes). **k**, Representative 200-gene H3K27ac-Pol II S5P dual-mark Epi-PHR and multiplexed FISH images. Magnified images display three decoded loci with their associated Epi-PHR signals in different Epi-PHR channels and multiplexed FISH signals in selected rounds of readout hybridization. Scale bar for the left images: 5 µm; scale bar for the magnified images: 500 nm. **l**, Stitched DAPI images of mouse brain hippocampus region. Analyzed CA (red boxes) and DG (blue boxes) fields of view are shown. Scale bar for the stitched image: 200 µm. **m**, H3K27ac and Pol II S5P Epi-PHR signal-to-background enrichment Z-scores of genes in CA and DG regions.

To determine whether the second set of hairpins functioned as effectively as the first set, we conducted an Epi-PHR experiment using the probe set that targets the 100-kb H3K9me3 enriched locus. Following the Epi-PHR procedure, we observed strong Epi-PHR signals colocalizing with FISH foci (Extended Data Fig. 3a), indicating that the second set of Epi-PHR hairpins performed as designed. Additionally, we performed systematic negative control experiments by omitting one key Epi-PHR component at a time (Extended Data Fig. 3a).

Consistent with the results from the first set of Epi-PHR hairpins, the omission of any Epi-PHR component eliminated the H3K9me3 Epi-PHR signals (Extended Data Fig. 3b). This demonstrated that the complete Epi-PHR assembly is required to generate specific Epi-PHR signals when using the second set of Epi-PHR hairpins.

To demonstrate dual-mark Epi-PHR, we used the 200-gene library to profile H3K36me3 and H3K27me3 simultaneously in the same single cells. In this experiment, we used the first set of hairpins for H3K36me3 and the second set for H3K27me3. From the raw images, we observed strong Epi-PHR signals from both channels and clear multiplexed FISH signals from each round of readout probe hybridization (Fig. 4b). After decoding loci identities and measuring Epi-PHR signal rates, we found that the dual-mark profiling results were highly reproducible, with Pearson correlation coefficients of 0.93 for H3K36me3 and 0.91 for H3K27me3 between biological replicates (Figs. 4c and 4d). Furthermore, the dual-mark profiling results strongly correlated and agreed with the single-mark profiling results (Figs. 4e and 4f), highlighting the reliability of the dual-mark system. Notably, when using the newly designed second hairpin set for H3K27me3 in dual-mark profiling, the Epi-PHR signal rates were strongly correlated with the result from the single-mark profiling of H3K27me3 using the first set of Epi-PHR hairpins (Fig. 4f). This added another line of evidence that the second set of Epi-PHR hairpins functioned as effectively as the first set.

To further test our dual-mark profiling approach, we cross-validated the dual-mark profiling results with ChIP-seq data^48^. For both H3K36me3 and H3K27me3 marks, we observed higher Epi-PHR signal rates in loci with high ChIP-seq peak densities of the same marks (Figs. 4g and 4h). Furthermore, Epi-PHR signal rates correlated strongly with ChIP-seq results of the same marks, with Pearson correlation coefficients of 0.80 for H3K36me3 and 0.65 for H3K27me3 (Fig. 4i and Extended Data Fig. 4a). These findings are consistent with our single-mark profiling experiments. Additionally, we calculated the probability of single mark enrichment and dual mark enrichment for each targeted gene (Fig. 4j). We observed a significantly higher probability of single mark enrichment, aligning with previous studies reporting that these two marks are largely mutually exclusive in gene bodies^1,48^. To further explore the possibility of dual-mark enrichment among the 200 selected genes, we examined genes with the highest probabilities of dual-mark enrichment in both replicates (Extended Data Figs. 4b and 4c). Intriguingly, we found that these genes were all close to TAD boundaries (Extended Data Figs. 4d and 4e), suggesting that the dual-mark enrichment may be related to TAD boundary dynamics and/or mechanics^58,59^. Finally, measuring the nuclear positions of mark-enriched loci revealed that H3K27me3-enriched loci were closer to the nuclear periphery (Extended Data Fig. 4f), consistent with our previous observations (Extended Data Fig. 2f). These analyses demonstrate the reliability of Epi-PHR system for dual-mark profiling in single cells.

We next sought to demonstrate two-mark histone profiling in mouse brain hippocampus tissue sections. For this experiment, we selected 200 genes including major cell type markers^60^ and subregion markers^61^ of the mouse hippocampus. We performed dual-mark Epi-PHR targeting H3K27ac and RNA Pol II S5P and identified genes by multiplexed FISH as above (Fig. 4k).

Based on Epi-PHR signals, we were able to identify distinct epigenetic states in manually segmented hippocampus cornu ammonis (CA) and dentate gyrus (DG) regions (Figs. 4l and 4m). Notably, CA and DG regions respectively showed active mark enrichment in genes known to be differentially expressed in major cell types within these regions (Fig. 4m; e.g. DG: *Glis3* and *Fam163b*; CA: *Lingo* and *Man1a*), based on scRNA-seq data^60^. The results here demonstrate that Epi-PHR can be applied to complex tissue and map spatial variations of epigenetic marks across tissue regions at the single-gene level.

### Epi-PHR can be combined with chromatin tracing to profile epigenetic state and chromatin folding at the same genomic target

To further demonstrate the multiplexing capabilities of Epi-PHR, we combined Epi-PHR with chromatin tracing^42^ to profile both the epigenetic state and 3D chromatin folding at the same genomic target in the same single cells. We selected as an Epi-PHR target a 500-kb region on human chromosome 20 known to be enriched with H3K27ac in IMR90 cells^48^. For chromatin tracing, we subdivided this 500-kb region into 30 equal (∼16.6-kb) tracing loci. FISH probes tiling this entire region were designed with a PH1 docking sequence, while chromatin tracing was facilitated by adding different readout sequences to FISH probes targeting the different tracing loci. During imaging, we first captured H3K27ac Epi-PHR signals from whole 500-kb regions, and then sequentially visualized the spatial positions of each tracing locus within the 500-kb target region (Extended Data Fig. 5a).

We next assembled 3D traces from sequential tracing images and measured Epi-PHR signals colocalized with each trace in Epi-PHR images (Extended Data Figs. 5b and 5c). As confirmation of preserved genome structure during the Epi-PHR protocol, chromatin tracing of the target region revealed structural features, including sub-TAD boundaries and loop anchors (Extended Data Fig. 5d). Critically, the structural patterns observed in our combined Epi-PHR and chromatin tracing dataset matched those from Hi-C^21^ (Extended Data Fig. 5e), validating the non-disruptive nature of Epi-PHR. Furthermore, inversed Hi-C^21^ contract frequency highly correlated with the median spatial distance between tracing loci, with a correlation coefficient of 0.86 (Extended Data Fig. 5e). Together, these results demonstrate the compatibility of Epi-PHR with chromatin tracing.

Before the present work, methods that reveal both high-resolution epigenetic mark profile and 3D chromatin folding conformation in real space were notably absent from the omics toolkit. However, epigenomic MERFISH^44^ is theoretically capable of profiling both epigenetic state and 3D chromatin folding in the same single cells. However, to the best of our knowledge, this capability has not yet been demonstrated. Therefore, to more comprehensively compare Epi-PHR to existing tools, we attempted the combination of epigenomic MERFISH with chromatin tracing. We used the same chromatin tracing probe library as above, targeting the 500-kb H3K27ac enriched region on chromosome 20. In this experiment, we first performed the epigenomic MERFISH protocol and imaged the sample for both epigenomic MERFISH and DAPI (whole nuclear DNA stain) signals. We then completed a standard chromatin tracing procedure on the same sample. We used DAPI signals to realign the sample and imaged the same cells as in the epigenomic MERFISH imaging step. Strong H3K27ac epigenomic MERFISH signals were generated from our genomic target, as were chromatin tracing DNA FISH signals (Extended Data Fig. 5f, left panel). However, chromatin tracing signals did not colocalize with epigenomic MERFISH signals. Furthermore, the chromatin tracing signals were so dispersed in the nuclei that no discernable structural features were preserved in assembled traces and the spatial distance matrix (Extended Data Fig. 5f, right panel), indicating severe disruption of chromatin architecture. We suspect this disruption resulted from the genomic fragmentation due to Tn5 tagmentation in epigenomic MERFISH, which reduces chromatin stability during the chromatin tracing DNA FISH procedure. As further evidence of this, we also observed global changes in the DAPI patterns between the epigenomic MERFISH and chromatin tracing steps (Extended Data Fig. 5f, left panel). Because tagmentation is inherent to epigenomic MERFISH, this challenge could be difficult to overcome. This demonstration further highlights the unique advantages of Epi-PHR for enabling combined high-resolution 3D genome tracing and locus-specific epigenetic profiling in the same single cells.

### Epi-PHR and chromatin tracing of an imprinted gene locus reveals allele specific epigenetic state and chromatin conformation

While most systems of gene expression in mammalian tissues utilize mechanisms of regulation that act equally on both copies of the genome within a diploid cell, epigenetic imprinting is a critical form of transcriptional regulation that selectively acts on genomic loci depending on their parent-of-origin^62–65^. Epigenetic imprinting is found in all metatherian and eutherian mammals and is theorized to be the outcome of the unequal selective pressures on maternal and paternal genes that arise from internal gestation and result in maternal/fetal conflict^66,67^. Most autosomally imprinted genes regulate metabolic and developmental pathways that have a direct impact on placental development, fetal size, and maternal/fetal metabolic regulation^67–75^. At the mechanistic level, gene expression regulation by epigenetic imprinting may involve critical parent-of-origin-specific chromatin folding^76,77^, making it desirable to obtain information about both epigenetic state and chromatin architecture at the same single copies of imprinted regions. However, such multiplexed and single-copy analysis is beyond the reach of current sequencing-based methodologies^21,78,79^. We reasoned that Epi-PHR combined with chromatin tracing, phased by FISH probes targeting single-nucleotide polymorphisms (SNPs)^80^, would overcome these limitations and allow for single-cell epigenetic modification quantification and chromatin folding with parent-of-origin specificity.

To demonstrate this capability, we leveraged Epi-PHR combined with chromatin tracing to study the association between epigenetic imprinting state and chromatin conformation at *Meg3*, a gene known to be paternally imprinted (maternally expressed) inside the *Dlk1-Meg3* locus^81^. As a model system, we utilized F1 hybrid CAST x C57 *in vitro* TSCs since trophoblastic tissues display well defined epigenetic imprinting and grow uniformly and robustly^71,82,83^. To validate the imprinting of *Meg3* in our TSC model, we performed RNA-seq on these cells and found the expected maternal bias of *Meg3* expression, as well as the expected allelic bias of other imprinted genes (Extended Data Figs. 6a and 6b). Known epigenetic mechanisms of imprinting in the murine model include canonical differential cytosine methylation (5mC) of imprinting control regions (also called differentially methylated regions, DMRs) and associated histone modifications H3K9me2/3^78,84–87^. Regulation of the *Dlk1-Meg3* locus is complex and involves three paternally methylated DMRs^81^, including the critical *Meg3-*DMR spanning the *Meg3* promoter and extending into its first intron^88^ (Fig. 5a). In addition to differential DNA methylation at the *Meg3*-DMR, the *Meg3* promoter has been shown to display paternal-specific enrichment of H3K9me3 histone modification^89^.

**Fig. 5.**
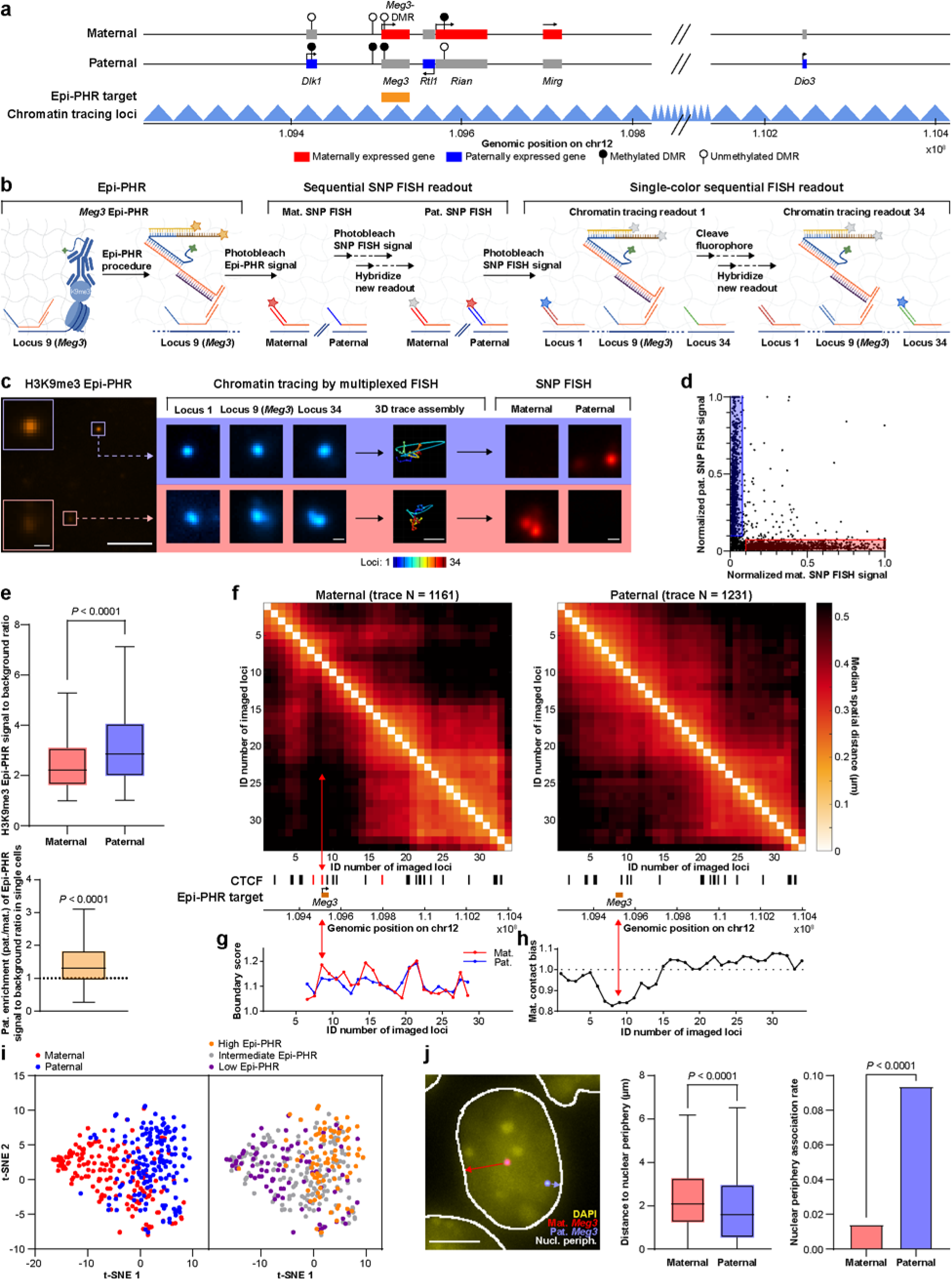
| Combined Epi-PHR and chromatin tracing of an imprinted locus in mouse trophoblast stem cells. **a**, Map of the *Dlk1-Meg3* imprinted locus and combined Epi-PHR and chromatin tracing experimental design. Epi-PHR probes tile the *Meg3* coding region, while FISH probes for tracing span 1.19 Mb in 34 loci (35-kb resolution). **b**, Experimental flow of Epi-PHR combined with chromatin tracing across the *Dlk1-Meg3* imprinted gene locus. PH1 hybridizes to FISH probes within the *Meg3* coding region via an adapter. During imaging, Epi-PHR signals were recorded and photobleached. Next, single nucleotide polymorphism (SNP) FISH signals were sequentially generated, imaged, and photobleached. Finally, tracing loci were visualized by 34 rounds of single-color imaging. **c**, Representative H3K9me3 Epi-PHR and chromatin tracing images. Sequential FISH foci are assembled into 3D chromatin traces. Traces are then assigned a parent-of-origin by spatial proximity to SNP FISH signals. Scale bars for original images: 5 µm; scale bars for magnified images: 500 nm. **d**, Populations of maternal and paternal traces are identified based on colocalization with normalized SNP FISH signals. **e**, (Top) H3K9me3 Epi-PHR signal-to-background is lower in the maternal *Meg3* than the paternal *Meg3*. Statistical significance was determined by an unpaired two-tailed Mann-Whitney test. Sample sizes: maternal *Meg3* (n = 1161), paternal *Meg3* (n = 1231). (Bottom) Single-cell paternal enrichments (paternal/material) of H3K9me3 Epi-PHR signal-to-background ratios are significantly above one, as determined by a one-sample Wilcoxon signed rank test. Sample size: n = 682 cells. In box plots, central lines represent median values, boxes show interquartile ranges (IQRs), and whiskers indicate non-outlier minimum and maximum values. Outliers were defined as values further than 1.5 times the IQR away from the box. **f**, (Left and right) Median inter-loci spatial distances of maternal and paternal *Dlk1-Meg3* chromatin traces. The red arrow highlights the location of the *Meg3*-DMR and a maternal-specific CTCF peak^90^. **g**, Boundary scores between tracing loci across the *Dlk1*-*Meg3* region. The red arrow highlights the location of the *Meg3*-DMR and a maternal-specific CTCF peak^90^. **h**, Maternal contact bias of each traced locus to all other tracing loci. The red arrow highlights the location of the *Meg3*-DMR. **i**, (Left) t-SNE of phased *Dlk1-Meg3* chromatin trace median spatial distances reveals parent-of-origin specific clustering. Traces were required to have 80% detection efficiency overall, and not more than two neighboring missing loci. Missing values were filled in by linear interpolation. Number of traces plotted for each parent-of-origin were equalized by random selection. Sample sizes: maternal traces (n = 167), paternal traces (n = 167). (Right) Relabeling the t-SNE plot at left according to H3K9me3 Epi-PHR level (high Epi-PHR = top 25% traces, intermediate Epi-PHR = middle 50% traces, low Epi-PHR = bottom 25% traces) reveals relationship between epigenetic state and 3D chromatin folding. Sample sizes: high Epi-PHR traces (n = 84), intermediate Epi-PHR traces (n = 166), low Epi-PHR traces (n = 84). **j**, Analysis of *Meg3* nuclear distribution relative to DAPI signal boundaries. (Left) Representative image of maternal and paternal *Meg3* FISH signals, DAPI, and nuclear periphery. (Middle) Paternal copies of *Meg3* are closer to the nuclear periphery than are maternal *Meg3* copies. Statistical significance was determined by an unpaired two-tailed Mann-Whitney test. The central lines represent median values, the boxes show interquartile ranges (IQRs), and whiskers indicate non-outlier minimum and maximum values. Outliers were defined as values further than 1.5 times the IQR away from the box. (Right) Paternal copies of *Meg3* are more frequently in association with the nuclear periphery than are maternal *Meg3* copies, as determined by a threshold distance of 200 nm. Statistical significance was determined by Fisher’s exact test. Sample sizes: maternal (n = 998), paternal (n = 1036).

Here, we performed H3K9me3 Epi-PHR with probes tiling the *Meg3* coding region (mm39 chr12: 109506880-109538160, Fig. 5a). To determine the chromatin folding around *Meg3*, we segmented an area covering the entire *Dlk1-Meg3* locus into 35-kb bins (34 bins starting at mm39 chr12:109226880, Fig. 5a), such that the entire *Meg3* was located inside of chromatin tracing target locus 9. We tiled each of these bins (except for the region covered by Epi-PHR probes) with chromatin tracing probes lacking PH1 docking sequences. Both Epi-PHR and chromatin tracing probes contained tracing-locus-specific readout sequences (Fig. 5b). To distinguish between maternal and paternal *Meg3* alleles, we additionally designed DNA SNP FISH probes targeting clusters of SNPs between the CAST and C57 genomes in proximity to the *Dlk1*-*Meg3* locus (mm39 chr12: 107000000-108000000 and 111000000-112000000). SNP FISH probes did not have PH1 docking sequences, but instead had parent-of-origin specific readout sequences for secondary FISH readout.

During imaging, we first captured Epi-PHR and DAPI signals. After photobleaching Epi-PHR signals, we sequentially imaged the SNP FISH signals for the two *Meg3* alleles. Finally, we sequentially imaged all tracing loci to determine the spatial positions of all genomic segments in the *Dlk1-Meg3* locus (Fig. 5b). After we assembled 3D chromatin traces, we identified colocalized SNP FISH signal for each trace, and were able to identify both maternal and paternal copies of *Meg3* (Figs. 5c and 5d). For each trace, we then calculated a H3K9me3 Epi-PHR signal-to-background ratio colocalized with the 3D position of the *Meg3* copy in that trace.

H3K9me3 Epi-PHR signals were higher for paternal *Meg3* alleles than in maternal alleles, both at the bulk and single-cell levels (Fig. 5e), in agreement with previous work^89^ and the expected imprinting status of the *Dlk1-Meg3* locus^81^. To validate this result, we performed H3K9me3 Cleavage Under Targets and Release Using Nuclease (CUT&RUN)^27^ in the TSCs and found an H3K9me3 bias to the paternal allele of *Meg3* (Extended Data Figs. 6c and 6d). Our CUT&RUN data yielded some H3K9me3 reads in the maternal allele of *Meg3* (Extended Data Fig. 6d), which is consistent with the lower but detectable H3K9me3 Epi-PHR signals of the maternal allele (Figs. 5c and 5e).

By analyzing our phased chromatin traces, we found allelic differences in the 3D chromatin folding across the *Dlk1-Meg3* locus (Fig. 5f). Most notably, we observed a maternal specific boundary at the 5’ edge of the *Meg3* locus (Figs. 5f and 5g), in agreement with a previous report that showed maternal specific CTCF binding at the *Meg3*-DMR and associated boundary formation^90^. We also observed increased maternal isolation (decreased maternal contact) of *Meg3* and nearby chromatin from the rest of the traced region (Fig. 5h). Dimensionality reduction of single-trace spatial distance matrices by *t*-Distributed Stochastic Neighbor Embedding (t-SNE)^91^ revealed distinct maternal and paternal structural clusters at the single trace level (Fig. 5i, left panel). In addition, grouping traces by H3K9me3 Epi-PHR signal levels revealed a similar pattern of trace clustering, with high Epi-PHR traces corresponding to the paternal traces and low PHR traces corresponding to the maternal traces (Fig. 5i), indicating that epigenetic state is associated with specific 3D chromatin conformations at the *Dlk1-Meg3* locus.

When we analyzed the spatial distribution of the *Meg3* loci inside the nucleus, we found that the inactive paternal copy of *Meg3* was generally closer to the nuclear periphery than the maternal copy (Fig. 5j, left and middle panels). Furthermore, the contact frequency to the nuclear periphery (contact defined as distance to nuclear periphery ≤ 200 nm) of the paternal allele was 9.4%, while that of the maternal allele was 1.4% (p < 0.0001, Fig. 5j, right panel). These findings are consistent with the known trend of repressed loci being localized to the nuclear periphery^92^, again demonstrating that the Epi-PHR procedure preserves both fine (chromatin folding structure) and large (sub-nuclear localization) scale genome organization.

Our Epi-PHR analyses above comparing the two parental origins support an expected, simple model from bulk allelic CUT&RUN and Hi-C analyses that H3K9me3 abundance is a surrogate of DNA methylation status at the *Meg3* locus: High H3K9me3 indicates methylated *Meg3* which blocks CTCF binding and in turn abolishes the domain boundary. We noticed that, although the paternal group overall showed higher H3K9me3 Epi-PHR signals compared to the maternal group, each group had a spectrum of different H3K9me3 levels and the distributions of the two groups overlapped significantly (Fig. 5e). We hypothesized that within the maternal group, chromatin traces with higher *Meg3* H3K9me3 signals may adopt paternal-like folding conformations without a domain boundary at *Meg3*, whereas within the paternal group chromatin traces with lower *Meg3* H3K9me3 signals may adopt maternal-like conformations with a domain boundary at *Meg3*. To test these hypotheses, we divided maternal and paternal traces into five quintile groups each based on their *Meg3* H3K9me3 levels. Surprisingly, *Meg3* boundaries among maternal traces became increasingly prominent with increasing H3K9me3 levels. Instead of adopting a paternal-like, boundary-less conformation, the highest H3K9me3 quintile maternal traces showed the most striking boundary at *Meg*3 (Fig. 6a, red arrows). Furthermore, paternal traces lacked a boundary at *Meg3*, even at low H3K9me3 levels (Figs. 6b, red arrows). For the paternal traces, decrease in H3K9me3 corresponded to an overall decompaction of the whole traced region, instead of an appearance of *Meg3* boundary or a maternal-like conformation (Fig. 6b). These observations enabled by the multi-omic single-copy profiling with Epi-PHR and chromatin tracing argue against the basal hypothesis from bulk allelic CUT&RUN and Hi-C analyses that H3K9me3 is a simple surrogate of DNA methylation which abolishes the *Meg3* boundary. Instead, maternal *Meg3* H3K9me3 level is positively associated with the *Meg3* boundary, and paternal *Meg3* H3K9me3 is associated with general chromatin compaction.

**Fig. 6.**
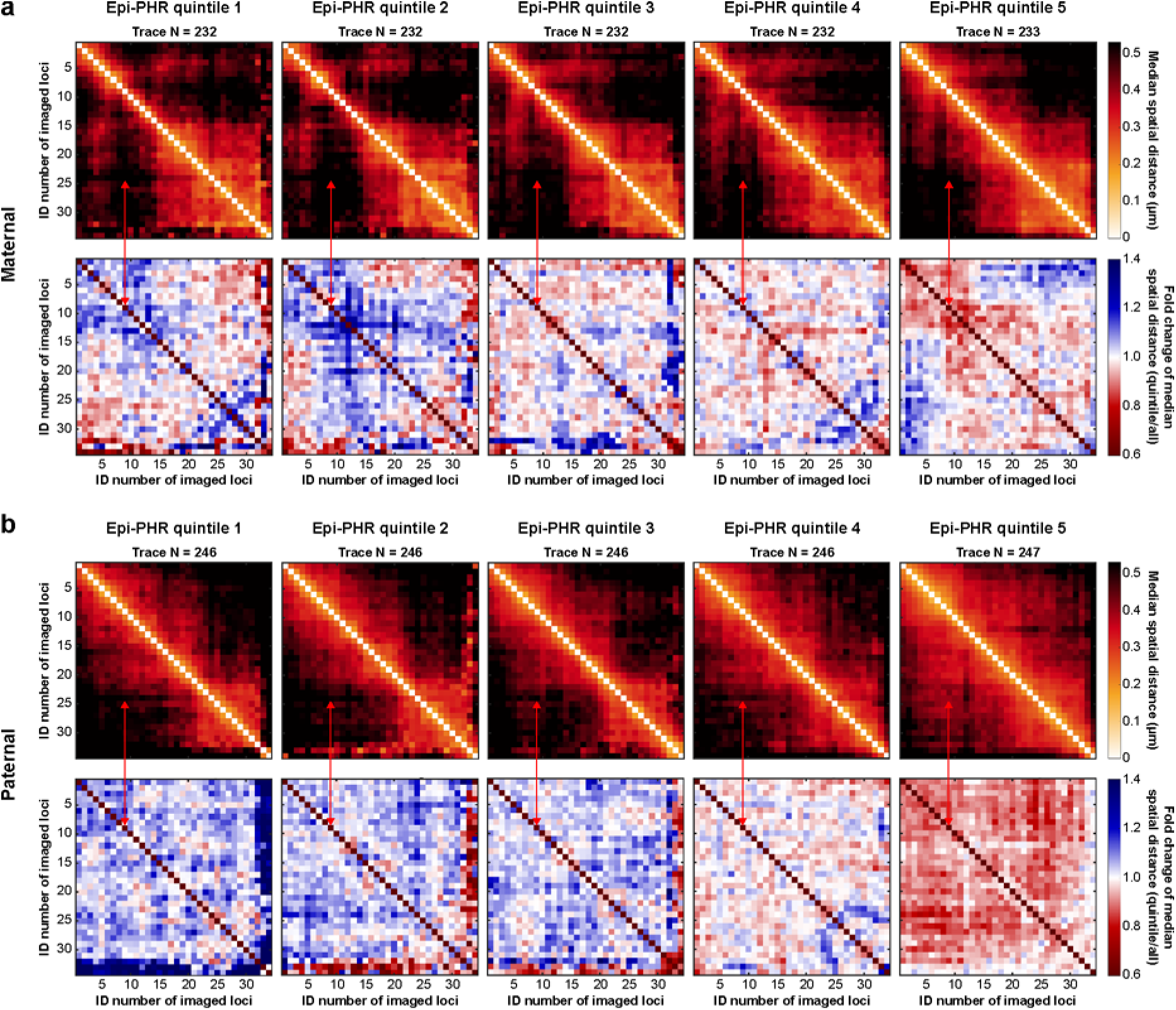
| Single-cell analysis of Epi-PHR and chromatin tracing of an imprinted locus in mouse trophoblast stem cells. a, Top: Median inter-loci spatial distance matrices of maternal traces divided into five quintile groups based on H3K9me3 Epi-PHR signal to background ratios. Quintiles 1-to-5 correspond to the lowest-to-highest H3K9me3 groups. The red arrows highlight the location of the *Meg3*-DMR. Bottom: Fold change of distance matrices of maternal trace quintile groups in comparison to all maternal traces. **b,** Top: Median inter-loci spatial distance matrices of paternal traces divided into five quintile groups based on H3K9me3 Epi-PHR signal to background ratios. Bottom: Fold change of distance matrices of paternal trace quintile groups in comparison to all paternal traces.

## Discussion

In this work, we introduced an epigenetic profiling method that can simultaneously profile more than one epigenetic mark at numerous genomic loci with single-gene or finer resolution, is non-disruptive and retains 3D chromatin architecture, and inherently provides single-cell and *in situ* information. Our method utilizes a system of DNA hairpins that are opened in the proximity between a genomic target and an epigenetic mark of interest. Signals representing this spatial proximity are visualized by fluorescence microscopy. We have shown that Epi-PHR is able to generate high signal-to-background signals even at genomic targets of 2-kb length, which potentially allows targeting of many short genes and other genomic elements. We also demonstrated the capability of Epi-PHR to profile various active and inactive histone modifications, as well as RNA Pol II. Because Epi-PHR only relies on proximity between an antibody and a genomic target, we anticipate Epi-PHR will allow profiling of numerous chromatin-associated features beyond histone modifications, such as transcription factors and DNA modifications. We also highlighted the true single-cell profiling capabilities of Epi-PHR, which offers an advantage over traditional bulk sequencing-based methods, as well as single-cell and spatial epigenetic mark sequencing methods, which are often limited in single-cell/single-spatial-spot reads, and may have more than one cells present in a given spatial spot.

Epi-PHR is readily compatible with other FISH-based methods for performance improvement and/or multi-omic information. By combining Epi-PHR with DNA MERFISH^49^, for example, we determined both the spatial positions of hundreds of genes and their epigenetic states in the same single cells in both human cell cultures and mouse brain tissue. By combining Epi-PHR with fine-scale chromatin tracing, we revealed the relationship between epigenetic state and chromatin folding at an imprinted locus. The chromatin tracing method is highly adaptable to a wide range of genomic resolutions to reveal various types of chromatin organization, and to all these Epi-PHR may be applied to add epigenetic information. Importantly, unlike epigenomic MERFISH, the Epi-PHR procedure preserves the 3D genome organization and allows for profiling of repressive epigenetic marks. Epi-PHR also expands to dual mark profiling, which helped us uncover an intriguing association between H3K36me3-H3K27me3 dual mark enrichment and the proximity to TAD boundaries. This association indicates that the higher tendencies for these regions to enrich both the active and inactive marks may be due to the dynamics and/or mechanics of chromatin folding through cohesin loop extrusion that establishes the TAD organization^58,59^. The advantages of Epi-PHR in comparison to other single-cell/spatial epigenetic mark profiling methods are summarized in Extended Data Fig. 7.

The Epi-PHR results presented here profiling the epigenetic state of *Meg3* represent the first phased single-cell examination of histone mark deposition at an imprinted gene. We combined this examination with chromatin tracing, additionally providing both single-cell and aggregate phased chromatin folding and nuclear localization information. Notably, we provided the first direct 3D trace visualization of the whole *Dlk1-Meg3* locus, confirming a maternal-specific chromatin domain boundary that was previously observed using bulk sequencing-based methods^90,93^. We further showed the close relationship between the chromatin folding of the *Dlk1-Meg3* locus and its epigenetic state at the single chromatin copy level. Surprisingly, among the maternal copies, the relationship between *Meg3* H3K9me3 level and the domain boundary is opposite to what one would expect, suggesting that H3K9me3 may enhance boundary formation at the maternal *Meg3* locus, though e.g. promoting chromatin interaction^94^. We note that such insight could not be achieved by integrating bulk allele-specific CUT&RUN and Hi-C analyses. Neither can it be achieved by a hypothetical, combined single-cell allele-specific CUT&TAG and Hi-C approach due to the limited read coverage and genomic resolution of single-cell CUT&TAG data (Extended Data Fig. 7), which would be even worse when sequencing reads without allelic SNPs need to be excluded. This unexpected finding is currently uniquely enabled by the single-copy, high-resolution, multi-omic capacity of Epi-PHR.

Epi-PHR phased by SNP FISH has several advantages over sequencing-based methods for studying allele-specific epigenetic state. For example, because phased Epi-PHR can rely on spatial proximity to a nearby region tiled with SNP FISH probes for phasing and not necessarily on SNPs in the Epi-PHR target locus itself, Epi-PHR combined with SNP FISH potentially allows measurements of challenging allelic targets that lack enough SNPs for phasing in traditional short-read-sequencing-based methods. Following a similar principle, FISH probes targeting adjacent sequence differences can be used to individually identify spatially distinct genomic targets with identical sequences, such as distant paralogs and pseudogenes, retrotransposons, and other dispersed repetitive elements. We expect this flexibility will allow researchers to distinguish among individual copies/clusters of many different classes of Epi-PHR targets that may contain identical sequences.

The non-disruptive nature and tissue-compatibility of Epi-PHR open many new avenues for studying epigenome, 3D genome, and gene regulation at the single-cell level in complex tissues. For example, Epi-PHR combined with fine-scale chromatin tracing readily allows for studying enhancer epigenetic states and 3D folding of *cis*-regulatory regions of genes for cell identity and cell state specification in heterogeneous tissue microenvironments. Multi-locus Epi-PHR can also help define a single-cell epigenetic aging clock or a single-cell epigenetic cancer biomarker by integrating the epigenetic states of hundreds of key gene regions, and do so in complex tissues with preserved spatial information. The dual-mark capacity of Epi-PHR will facilitate studies of mechanisms underlying dual mark deposition and maintenance on the same copy of gene, which is not permitted by state-of-the-art techniques such as Spatial-CUT&Tag^40^ (which does not have single-copy resolution) and epigenomic MERFISH^44^ (which cannot profile two marks). In sum, Epi-PHR gives access to the world of spatial epigenetic heterogeneity that has to this point been challenging to explore.

## Methods

### Library design

All library, primer, and readout sequences can be found in Supplementary Table 1.

#### Genomic-resolution tests

We selected a 3.3-Mb region spanning hg19 chr1:3300001-6600000 that was previously shown to contain high levels of H3K9me3 in IMR90 cells^48^ and that we also noted showed relatively homogeneous 3D structure across its length^21^. We split this sequence into 1-kb fragments and then processed these fragments with OligoArray2.1^95^ using the following parameters: Oligos must be 30 nt in length, and may not overlap; the melting temperature must be between 65°C and 85°C; the GC content must be between 30% and 90%; there can be no secondary structure with a melting temperature greater than 70°C; there can be no cross-hybridization among oligos with a melting temperature greater than 70°C; single-nucleotide repeats of length 6 or more were not allowed. At this initial design step, OligoArray2.1 was provided with FASTA database for its built-in specificity filtering step consisting of a single sequence of ∼70 kb repeated “A” nucleotides. Oligos were then filtered for binding specificity in two steps: (1) by screening against a database of repetitive sequences in the human genome, downloaded from Repbase^96^ [https://www.girinst.org/repbase/, Repeat class: “All”, Taxon: “Homo sapiens (Human)”, including ancestral] using BLAST+ blastn-short^97^, allowing no match score equal to or greater than 33 to remove any oligo with alignments of 17 consecutive bases or more, and (2) by screening against the unmasked human genome (hg19) using BLAST+ megablast^98^ and requiring exactly one match per oligo. Once specific oligos tiling the whole 3.3-Mb region were generated, we then integrated the epigenetic profile^48^ and distribution of designed oligos to select six overlapping regions of increasing length (2 kb, 5 kb, 10 kb, 20 kb, 50 kb, and 100 kb) while balancing the following priorities: maximizing the overall level of H3K9me3 in selected regions, ensuring even scaling of total H3K9me3 levels across resolutions, and ensuring even scaling of number of tiling oligos across resolutions. Based on these considerations, we selected a 5’ start coordinate for all resolutions at hg19 chr1:4742385. For each resolution we assembled encoding and docking sequences in the following configuration (5’ to 3’): a 30-nt PHR docking sequence, a 30-nt readout sequence, and the 30-nt encoding sequence generated above (docking and readout sequences were taken from^42^). Finally, 20-nt primers were added to each end of all assembled oligos. Because the sequences of each oligo are reverse complemented during Library Synthesis (see below), the internal sequences were reverse complemented (from the intended final probe sequences) before adding primers. Oligo pool libraries were synthesized by Twist Bioscience.

#### 200-gene single- and dual-mark Epi-PHR

We selected 200 genes with lengths ranging from 25 kb to 35 kb (the gene list is provided in Supplementary Table 2). The gene body sequences were downloaded from Roadmap Epigenomics Consortium database using hg19 assembly. We processed these sequences with ProbeDealer^99^ using the following parameters: Oligos must be 30 nt in length, and alternate among plus and minus strand with 5 nt overlaps; the melting temperature must be between 65°C and 85°C; the GC content must be between 33% and 83%; single-nucleotide repeats of length 6 or more were not allowed. Oligos were then filtered for binding specificity in two steps: (1) by screening against a database of repetitive sequences in the human genome, downloaded from Repbase^96^ [https://www.girinst.org/repbase/, Repeat class: “All”, Taxon: “Homo sapiens (Human)”, including ancestral] using BLAST+ blastn-short^97^, allowing no match with a score equal to or greater than 33 to remove any oligo with matches of 17 consecutive bases or more, and (2) by screening against the unmasked human genome (hg19) using BLAST+ megablast^98^ and requiring exactly one match per oligo. Next, we assigned a 36-bit, Hamming distance 2 (HD2) barcode to each targeted locus. To avoid signal crowing during the imaging, we required that each positive bit was only found once in a sliding window of 5 consecutive loci along each chromosome (the codebook is provided in Supplementary Table 2). Next, we assembled encoding and docking sequences in the following configuration (5’ to 3’): a 20-nt bit-specific readout sequence (derived from^49^), the 30-nt encoding sequence generated above, another copy of the 20-nt bit-specific readout sequence, and a 30-nt PHR docking sequence^42^. Finally, 20-nt primers were added to each end of all assembled oligos. Oligo pool libraries were synthesized by Twist Bioscience.

We also designed the genomic fiducial oligos for draft corrections among different rounds of multiplexed FISH hybridization. Chromosome-specific repetitive sequences were extracted from^100^, and highly conserved repeats with length longer than 26 nt were selected as genomic fiducial oligos. The detailed parameters for each targeted repeat are as follows: (hg19) chr1:2581275-2634211 contains 104 copies of targeted sequence, chr3:195199022-195233876 contains 200 copies of targeted sequence, and chrX:76096-94831, contains 71 copies of targeted sequence. All oligos contain 2 copies of 30-nt readout binding sequence and were ordered from Integrated DNA Technologies (IDT).

#### Epi-PHR in mouse hippocampus tissue

We selected 200 genes for Epi-PHR profiling in mouse hippocampus, utilizing publicly available scRNA-seq datasets^60^. To aid gene selection, we used a custom algorithm as well as NS-Forest^101^. Specifically, our custom algorithm involved taking the geometric mean of gene expression levels to pseudo-bulk the data for each major cell type, and then ranking candidate genes based on, 1) expression fold change relative to the mean expression level in all major cell types, 2) proportion of non-mitochondrial total gene expression within that cell type, and 3) gene length. Cell types with fewer than 40 cells within the mouse hippocampus scRNA-seq datasets were removed from the calculations. After an initial set of candidate genes was identified, we hand-checked genes against ISH (*in situ* hybridization) images available on the Allan Brain Atlas website (https://portal.brain-map.org/). From the candidate gene list, we selected a final set of 200 genes, including at least 15 genes for each of 12 major cell types, gene markers for different hippocampus sub-regions^61^, and 2 genes that were highly expressed across all 12 cell types based on the scRNA-seq data. We ensured that all selected genes were spaced at least 1 Mb apart within the genome, to limit optical crowding. After the final list of genes was selected, we designed oligos binding the antisense strands of the unspliced gene sequences to a maximum length of 20 kb from their transcription start sites using OligoArray2.1^95^ and the parameters described above for the genomic-resolution test. Source gene sequences were downloaded from the Ensembl website^102^. Oligos were then filtered for specificity using the approach described above for the genomic-resolution test. For this library, specificity filtering databases consisted of the repetitive sequences in the mouse genome, downloaded from Repbase^96^ [https://www.girinst.org/repbase/, Repeat class: “All”, Taxon: “Mus musculus (Mouse)”, including ancestral], and the unmasked mouse genome (mm39). We then assigned barcodes to genes and assembled the final library as described above for the 200-gene single- and dual-mark multiplexed FISH experiment. The final list of 200 genes and the complete codebook may be found in Supplementary Table 3.

We also designed the genomic fiducial oligos for drift corrections among different rounds of multiplexed FISH hybridization. Specifically, we targeted three 100 kb genomic regions, ensuring non-overlap with our gene targets (mm39; chr1:100500000-1005999999, chr12:100000000-100099999, and chr13:81200000-81299999). Oligos were designed using OligoArray2.1^95^ and the parameters described above for the genomic-resolution test. After specificity filtering as described above for the 200-gene mouse hippocampus library, we selected 333 oligos for each locus, appended two fiducial 20-nt readout sequences (derived from^49^), and finally 20-nt primer sequences at each end. Oligo pool libraries were synthesized by Twist Bioscience.

#### Epi-PHR with chromatin tracing in cell culture

We selected a 500-kb region spanning hg19 chr20:48700000-49200000 that was previously shown to be modified with H3K27ac in IMR90 cells^48^. This region was split into 30 equal tracing loci (∼16.6 kb). Encoding sequences were generated for each locus by further splitting each locus sequence into fragments of 1 kb (or the remainder), and then processing these sequences with OligoArray2.1^95^ using the following parameters: Oligos must be 30 nt in length, and may overlap up to 10 nt; the melting temperature must be between 65°C and 85°C; the GC content must be between 30% and 90%; there can be no secondary structure with a melting temperature greater than 70°C; there can be no cross-hybridization among oligos with a melting temperature greater than 70°C; single-nucleotide repeats of length 6 or more were not allowed. At this initial design step, OligoArray2.1 was provided with FASTA database for its built-in specificity filtering step consisting of a single sequence of ∼70 kb repeated “A” nucleotides.

Oligos were then filtered for binding specificity in two steps: (1) by screening against the unmasked human genome (hg38) using BLAST+ megablast^98^ and allowing only one match per oligo, and (2) by screening against a database of repetitive sequences in the human genome, downloaded from Repbase^96^ [https://www.girinst.org/repbase/, Repeat class: “All”, Taxon: “Homo sapiens (Human)”, including ancestral], using BLAST+ blastn-short^97^, allowing no match with a score equal to or greater than 33 to remove any oligo with matches of 17 consecutive bases or more. Next, 200 filtered oligos were selected for each tracing locus by utilizing the following algorithm until the number of required oligos was reached: (1) non-overlapping oligos were selected in a 5’ to 3’ direction; (2) overlapping oligos were sequentially added such that the degree of overlap between each current candidate oligo and already selected oligos was minimized; (3) oligos were duplicated, with the duplication spread out evenly across all previously selected oligos. Next, we assembled encoding and docking sequences in the following configuration (5’ to 3’): a 20-nt bin-specific readout sequence, the 30-nt encoding sequence generated above, an additional copy of the 20-nt bin-specific readout sequence, and a 30-nt PHR docking sequence (readout sequences were derived from^49^, while the PHR docking sequence was taken from^42^). Finally, 20-nt primers were added to each end of all assembled oligos. Because the sequences of each oligo are reverse complemented during Library Synthesis (see below), the internal sequences were reverse completed (from the intended final probe sequences) before adding primers. Oligo pool libraries were synthesized by Twist Bioscience.

##### Epi-PHR plus chromatin tracing of the Dlk1-Meg3 locus in mouse trophoblast stem cells

First, we selected a 1.2-Mb region spanning mm39 chr12:109218775-110418774. This sequence was next split into 1-kb fragments and then processed with OligoArray2.1^95^ using the following parameters: Oligos must be 30 nt in length, and may overlap by up to 5 nt; the melting temperature must be between 65°C and 85°C; the GC content must be between 30% and 90%; there can be no secondary structure with a melting temperature greater than 70°C; there can be no cross-hybridization among oligos with a melting temperature greater than 70°C; single-nucleotide repeats of length 6 or more were not allowed. At this initial design step, OligoArray2.1 was provided with FASTA database for its built-in specificity filtering step consisting of a single sequence of ∼70 kb repeated “A” nucleotides. Oligos were then filtered for binding specificity in two steps: (1) by screening against a database of repetitive sequences in the mouse genome, downloaded from Repbase^96^ [https://www.girinst.org/repbase/, Repeat class: “All”, Taxon: “Mus musculus (Mouse)”, including ancestral] using BLAST+ blastn-short^97^, allowing no match with a score equal to or greater than 33 to remove any oligo with matches of

#### 17 consecutive bases or more, and (2) by screening against the unmasked mouse genome (mm39) using BLAST+ megablast^98^ and requiring exactly one match per oligo

Next, specifically binding oligos were divided into 35-kb bins, starting at mm39 chr12:109226880, so that the imprinted gene *Meg3* was located entirely within the 9^th^ bin, and the range of bins spanned the area of known inter-allele variation in 3D genome structure^90^.

Within each bin, every other oligo was reverse complemented, and then oligos were duplicated, with the duplication spread out evenly across all oligos, until the total number was 1134, equal to the number of unique oligos in the bin with the highest number of unduplicated oligos. Next, we assembled encoding and docking sequences in the following configuration (5’ to 3’): a 20-nt bin-specific readout sequence, the 30-nt encoding sequence generated above, and two additional copies of the 20-nt bin-specific readout sequence (readout sequences were derived from^49^). For oligos binding within the *Meg3* gene body (mm39 chr12: 109506880-109538160), an additional the 30-nt PHR docking sequence (taken from^42^) was to the 5’ end of the assembly. Finally, 20-nt primers were added to each end of all assembled oligos. Because the sequences of each oligo are reverse complemented during Library Synthesis (see below), the internal sequences were reverse completed (from the intended final probe sequences) before adding primers. Oligo pool libraries were synthesized by Agilent (SurePrint Oligo Pool).

To distinguish the two alleles of *Meg3* in our hybrid mouse trophoblast stem cell (TSC) line, we designed a homologue-specific OligoPaints (HOPs) library^80^. We first extracted two 1-Mbp regions upstream (mm39 chr12: 107000000-108000000) and downstream (mm39 chr12: 111000000-112000000) of the *Meg3* cluster from the C57 and CAST genomes separately. These sequences were processed with ProbeDealer^99^ using the following parameters: oligos must be 30 nt in length and non-overlapping; the melting temperature must be between 65°C and 85°C; the GC content must be between 33% and 83%; and single-nucleotide repeats of length 6 or more were not allowed. Oligos were then filtered for binding specificity in three steps: (1) screening against the associated genome (C57 probes against the C57 genome and CAST probes against the CAST genome) using BLAST+ megablast^98^, requiring exactly one match per oligo; (2) screening against the opposite genome (C57 probes against the CAST genome and CAST probes against the C57 genome) using BLAST+ megablast^98^, requiring no match in the opposite genome; and (3) ensuring that two probes from different genome assemblies have more than 2 single nucleotide polymorphisms (SNPs) (See SNP identification details below). Next, we assembled encoding and readout sequences in the following configuration (5’ to 3’): two copies of 20-nt allele-specific readout sequence (derived from^103^), the 30-nt encoding sequence generated above, and two additional copies of the 20-nt allele-specific readout sequence. Finally, 20-nt primers were added to each end of all assembled oligos. Oligo pool libraries were synthesized by Twist Bioscience.

##### Estimation of spatial proximity distance threshold of Epi-PHR

We estimate the spatial proximity distance threshold of our Epi-PHR system at 60 nm, based on the combined length of the activator binding region, the DNA linker sequence and the PH1 docking adapter length on our Epi-PHR assembly (180 nt total). Assuming fully extended double-stranded DNA after PH1 docking and activator binding, the physical searching distance should be maximally 60 nm. In reality, most of the time our assembly with double-stranded and single stranded DNA regions would not be fully extended in solution, thus the 60 nm should be viewed as an upper limit.

##### Library synthesis

We synthesized batches of primary oligos from template libraries using a previously published protocol^104^. In brief, template libraries were first amplified by limited-cycle PCR, using pairs of 20-nt PCR primers that bind at the ends of the template sequences. The forward primer contained an additional T7 promoter sequence at its 5’ end. This promoter sequence was then used during *in vitro* transcription to further amplify the library. Next, final DNA oligos were generated by reverse transcription of the RNA product. Finally, the DNA oligos were purified and concentrated by hydrolysis of the RNA product, column purification, and drying in a DNA SpeedVac vacuum concentrator (Thermo Fisher Scientific, DNA120-115). Primers were ordered from IDT. Primer sequences are provided in Supplementary Table 1.

##### Coverslip preparation

Glass coverslips (40-mm-diameter, #1.5, Bioptechs, 40-1313-03193) were first washed by submersion in 50:50 37% hydrochloric acid and methanol for 30 min at room temperature (RT), rinsed 3 times in Milli-Q water, rinsed once in 70% EtOH, and dried in a drying oven at 70°C. Coverslips were then silanized by submersion in chloroform with 0.1% v/v triethylamine (MilliporeSigma, TX1200) and 0.4% v/v allyltrichlorosilane (MilliporeSigma, 107778) for 30 min at RT, rinsed with chloroform once, rinsed with ethanol once, and dried in a drying oven at 70°C for 1 hr. Silanized coverslips were stored in a desiccator until needed.

##### Cell culture

###### IMR90

IMR-90 cells (ATCC CCL-186) were cultured following standard procedures (5% CO_2_, 37°C) in Eagle’s minimum essential medium (EMEM; ATCC, 30-2003) with 10% v/v fetal bovine serum (FBS; Avantor, 89510-186) and Penicillin-Streptomycin (Gibco, 15140-122, 1:100), on silanized coverslips. Cells were grown to no more than ten passages.

###### Trophoblast Stem Cells

Murine CAST Il x C57 Il F1 TSCs (gift of Mauro Calabrese) were cultured as previously described on gelatin coated plates pre-plated with irradiated CF1 murine embryonic fibroblasts (ATCC, SCRC-1040) in TSC media (RPMI 1640 [Gibco, 11875], 20% FBS [Gemini BenchMark, 100-106], 0.1 mM Penicillin-streptomycin [Gibco, 15140], 1 mM sodium pyruvate [Gibco, 11360], 1x GlutaMAX [Gibco, 35050], 100 µM β-mercaptoethanol [SigmaAldrich, M3148], 25 ng/mL FGF4 [SigmaAldrich, F8424], 1 µg/mL Heparin [SigmaAldrich, H3149])^71,82,83^ and media was exchanged every 24-48 hours. Cells were passaged by with incubation with 0.25% Trypsin-EDTA (Gibco, 25200), and disrupted by pipetting before re-plating. Colonies were passaged prior to confluency to prevent differentiation.

To generate cells on imaging slides, silanized coverslips were placed in empty culture plates and irradiated MEF’s were added directly onto the coverslips without prior gelatin coating. TSCs were cultured as above for 2-4 days, and visually inspected for colony growth. Samples observed to have spontaneously differentiated were discarded. All tissue culture was performed at 37°C and 5% CO_2_.

To remove MEFs for downstream sequencing-based analysis, passaged cells were plated to a new cell culture dish (of equivalent size to the prior dish) without prior gelatin-coating for 1 hr. Non-adherent cells were collected for downstream sequencing analysis.

##### Mouse Tissue

Animal care and experiments were carried out in accordance with NIH guidelines and were approved by the Institutional Animal Care and Use Committee of Yale University. At 8 weeks of age, female C57BL/6 mice were anesthetized with isoflurane and euthanized. Whole brains were collected and embedded in O.C.T. compound (Sakura, 4583) by rapid freezing on dry ice, and were then stored at -80°C for long-term preservation.

##### Tissue Sectioning

Tissue blocks were balanced in a cryostat (Leica, CM3050 S) for 30 min at -20°C, and then sectioned at 10 µm onto silanized coverslips additionally treated with poly-D-lysine (1 mg/mL; EMD Millipore, A-003-E) for 30 min at RT.

##### Epi-PHR

###### Sample Preparation

Samples were rinsed with Dulbecco’s phosphate-buffered saline (DPBS; Sigma-Aldrich, D8537), and were then fixed with 4% paraformaldehyde (PFA; Electron Microscopy Sciences, 15710) in DPBS for 10 min, and washed with DPBS for 5 min 3 times. Samples were permeabilized using either 0.5% v/v or 1.0% v/v Triton X-100 (Sigma-Aldrich, X100) in DPBS for 10 min at RT with gentle shaking, followed by washing with DPBS for 3 min 3 times. Samples were then treated with 0.1 M hydrochloric acid for 5 min at RT and washed with DPBS for 5 min 3 times. Samples were then digested with RNase (Invitrogen, AM2286; 1:50 or 1:100) in DPBS at 37°C for 45 min, and then rinsed with DPBS 3 times.

For work in mouse hippocampus tissue, sample preparation was performed as above, except that permeabilization was performed using 8% v/v Triton X-100 in DPBS for 30 min.

For work in TSCs, sample preparation was performed as above, except that permeabilization was performed using 0.5% v/v Triton X-100 in DPBS for 20 min.

###### Antibody Staining and Gel Embedding

Samples were then treated with an Endogenous Biotin-Blocking Kit (Invitrogen, E21390) following the manufacturer’s protocol. The samples were next incubated in blocking buffer consisting of 10% v/v Donkey Serum (MilliporeSigma, S30-M), 10% w/v Bovine Serum Albumin (Sigma-Aldrich, A9647), 300 mM glycine (AmericanBio, AB00730), and 0.1% v/v Tween-20 (Sigma, P7949) in DPBS for 30 min at RT. Samples were then incubated in primary antibody in blocking buffer for 1 hr at RT. Primary antibodies and working concentrations used were: anti-H3K9me3 donkey (Abcam, ab8898, 1:100), anti-H3K27ac (Abcam, ab4729, 1:100), anti-H3K36me3 (Abcam, ab9050, 1:100; Active Motif, 61022, 1:100), anti-H3K27me3 (Cell Signaling, C36B11, 1:20), anti-Pol II S2P (Abcam, ab5095, 1:50) and anti-Pol II S5P (Abcam, ab193467, 1:100), rabbit IgG Control (EpiCypher 13-0042, 1:50), mouse IgG control (Sigma-Aldrich 12-371, 1:100). After primary antibody staining, samples were washed with DPBS 3 times, and then incubated with secondary antibody in blocking buffer for 1 hr at RT, followed by washing with DPBS for 5 min 3 times. Secondary antibodies and working concentrations used were the following: biotin-conjugated donkey anti-rabbit (Jackson ImmunoResearch, 711-065-152, 1:1000), and unconjugated donkey anti-mouse (Jackson ImmunoResearch, 715-005-150, 1:1000). After antibody staining, samples were then fixed in 4% PFA in DPBS for 5 min at RT and washed with DPBS for 2 min 3 times. Samples were further fixed with 1.5 mM BS(PEG)9 (Thermo Scientific, 21582) in DPBS for 20 min at RT, washed with DPBS for 2 min 2 times, and quenched with 100 mM Tris PH 7.4 (AmericanBio, AB14044) for 10 min at RT. Samples were then embedded in 4% Acrylamide/Bis 19:1 gel (Bio-Rad, 1610144) for 1.5-2.5 hr at RT.

For work in TSCs, antibody staining was performed as above, except that primary antibodies were incubated on the sample overnight (>16hr) at 4°C.

For work in mouse hippocampus tissue, antibody staining was performed as in TSC experiments described above.

###### Primary FISH Hybridization

Samples were incubated at RT for 30 min in prehybridization buffer, consisting of 50% formamide (Sigma-Aldrich, F7503) and 0.1% v/v Tween-20 in 2xSSC. After prehybridization, primary hybridization buffer containing required oligo pool libraries was placed on the bottom of small cell culture dishes and the samples were inverted into the dishes to sandwich this buffer.

Primary hybridization buffer contained 50% formamide (Sigma-Aldrich, F7503) and 20% Dextran Sulfate (EMD Millipore, S4030) in 2xSSC (Invitrogen, 15557044). Oligo pool usage details are given for each experiment below. All primary hybridization reactions were performed in 100 µl volume. Samples were denatured in a water bath set to 83°C for 10 min, and then incubated at 37°C in a humid chamber for at least 16 hr. Following primary FISH incubation, samples were washed with 0.1% v/v Tween-20 in 2xSSC for 15 min 2 times in a 60°C water bath and 1 time at RT.

For work in mouse hippocampus tissue, primary FISH hybridization was performed as for cell culture, with the following exception: Primary FISH incubation was extended to 2 overnights (>36 hr).

For work in TSCs, primary FISH hybridization was performed as for cell culture, with the following exceptions: Primary FISH incubation was extended to 2 overnights (>36 hr). After primary FISH incubation, samples were washed with 60% v/v formamide in 2xSSC with 0.1% v/v Tween-20 for 15 min 2 times with gentle rocking at RT, followed by 1 wash in 0.1% v/v Tween-20 in 2xSSC for 15min at RT.

For single-molecule FISH Epi-PHR experiments, ∼650 ng of Epi-PHR library were used for each sample. For 200-loci single- and dual-mark Epi-PHR experiments, ∼750 ng genomic fiducial oligos were mixed with ∼20,000 ng 200-loci Epi-PHR library for each sample. For each mouse hippocampus Epi-PHR sample, ∼50,000 ng of 200-gene Epi-PHR library were mixed with ∼1,000 ng genomic fiducial oligos. For Epi-PHR with chromatin tracing in cell culture, ∼7,500 ng genomic fiducial oligos were mixed with ∼22,000 ng Epi-PHR with chromatin tracing library for each sample. For Epi-PHR plus chromatin tracing the *Dlk1-Meg3* locus in mouse trophoblast stem cells, ∼4,800 ng SNP FISH probes were mixed with ∼4,000 ng Epi-PHR probes targeting *Meg3* and ∼13,000 ng *Dlk1-Meg3* locus chromatin tracing probes for each sample.

###### Proximity Hybridization Reaction and Clearing

For single-mark Epi-PHR, hairpin sequences of PHR components PH1, PH2, H1, and H2 were sourced from^45^, with additional sequences added as needed (sequences are provided in Supplementary Table 1). Before use, PHR components were freshly refolded in HCR buffer consisting of 50 mM Na_2_HPO_4_ (Avantor, 3828-01) and 1 M NaCl (Sigma-Aldrich, S5150), adjusted to pH 7.4 with HCl, according to the following program in a PCR machine: 95°C for 10 min, -1°C temperature change every 1 min (72x), then hold at 4°C forever. Refolding was performed with 40 nM oligo concentration in 100 µL HCR buffer.

In the following steps, all low-volume incubation steps (<1 mL volume) were performed by applying the component mixture to parafilm in the bottom of a small cell culture dish, inverting the sample on top of the buffer, end enclosing the small dish in a larger cell culture dish with wet paper to create a humid chamber.

First, linker oligo was hybridized to samples at 40 nM concentration in 100 µL buffer (50% formamide and 20% dextran sulfate in 2xSSC) for 45 min at 37°C and were then washed in 2xSSC with 0.1% v/v Tween-20 in a 60°C water bath for 5 min 3 times. Next, samples were incubated with PH1 hybridization mixture (100 µL refolded PH1, 200 µL formamide, and 100 µL 50% Dextran Sulfate) for 45 min at 37°C and were then washed in 2xSSC with 0.1% v/v Tween-20 in a 60°C water bath for 5min 3 times. Samples were then incubated with 3 µg streptavidin (Jackson ImmunoResearch, 016-540-084; Jackson ImmunoResearch, 016-000-114) in 3 mL blocking buffer for 30 min at 37°C and were then washed in DPBS for 5 min 3 times. The samples were next incubated with PH2 hybridization mixture (100 µL refolded PH1, 35 µL formamide) for 45 min at 37°C, and were then washed in 35% formamide in 2xSSC with 0.1% v/v Tween-20 for 5 min 3 times. The activator oligo was hybridized to the sample at 40 nM concentration in 100 µL buffer (25% formamide, 20% Dextran Sulfate, in 2xSSC) for 1 hr at 37°C, and then washed with 35% formamide in 2xSSC with 0.1% v/v Tween-20 for 5 min 3 times. Samples were incubated with H1 hybridization mixture (100 µL refolded H1, 80 µL 5 M NaCl, 120 µL Dextran Sulfate, and 100 µL water) for 1 hr at 37°C and washed with 35% formamide in 2xSSC with 0.1% v/v Tween-20 for 5 min 3 times. H2 hybridization and washing were performed as for H1. Finally, blocking oligo was hybridized to samples at 100 nM concentration in 400 µL buffer (100 µL HCR buffer, 80 µL 5 M NaCl, 120 µL Dextran Sulfate, and 100 µL water) for 1 hr at 37°C and were then washed with 35% formamide in 2xSSC with 0.1% v/v Tween-20 for 5 min 3 times.

Immediately prior to imaging, samples were digested and cleared for 1 hr at 37°C in digestion buffer consisting of Proteinase K Solution (Invitrogen, 25530049, 1:50) and 2% SDS (Sigma-Aldrich, 05030) in 2xSSC with 0.5% Triton X-100 and were then washed in DPBS at minimum 10 min 3 times with gentle shaking. Excess gel was trimmed as needed to facilitate assembly into the imaging chamber and prevent flow channel blocking during imaging.

#### 200-gene single-mark Epi-PHR

For multiplexed Epi-PHR experiments, the general experiment protocol followed the procedure described above, with exceptions that the oligo conjugated secondary antibody were used for PH2 docking. The oYo-link oligo conjugation kit (alphaThera, AT1002-100ss) was used to prepare the oligo-conjugated antibody, following the manufacturer’s instructions. In brief, 5 μg secondary antibody (Jackson ImmunoResearch, donkey anti-mouse 715-005-150 and donkey anti-rabbit 715-005-152) was mixed with 5 μl oYo-link reagent, and placed under UV lamp for 2∼3 hours. To avoid non-specific binding of oligo conjugated antibody, we diluted 2 μl oligo conjugated antibody into 100 μl high salt blocking buffer consisting of 1 M NaCl, 20 mM HEPES, 10% w/v Bovine Serum Albumin, 300 mM glycine, 0.1% v/v dextran sulfate and 0.3% v/v Triton-X^43,105^. Then oligo conjugated secondary antibodies were incubated on the sample overnight (>16 hr) at 4°C, followed by washing with high salt blocking buffer for 5 min 3 times. To dock PH2 to oligo conjugated antibody, PH2 docking linker oligo and PH1 docking linker oligo were mixed and hybridized to samples at 40 nM concentration for each linker oligo in 100 µL buffer (50% formamide and 20% dextran sulfate in 2xSSC) for 45 min at 37°C and were then washed in 2xSSC with 0.1% v/v Tween-20 in a 60°C water bath for 5 min 3 times. After PH1 hybridization, the samples were incubated with PH2 hybridization mixture (100 µL refolded PH2, 100 µL formamide, 40 µL 20xSSC, 100 µL dextran sulfate, 60 µL water) for 45 min at 37°C, and were then washed in 35% formamide in 2xSSC with 0.1% v/v Tween-20 for 5 min 3 times. The rest of the procedure followed standard Epi-PHR protocol.

#### 200-gene dual-mark Epi-PHR

For dual-mark Epi-PHR, a second set of PHR hairpins was designed as described in^45^. Hairpin conformations and binding kinetics were calculated using NUPACK^106^ to ensure two sets of Epi-PHR hairpins had similar parameters and minimized crosstalk among two sets of Epi-PHR hairpins.

Following the antibody blocking step, 0.5 µg of H3K27me3 and 1 µg of H3K36me3 primary antibodies were mixed in 100 µL of blocking buffer and incubated for 1 hour at RT. After primary antibody staining, the samples were washed with DPBS three times and then incubated with oligo-conjugated anti-mouse and anti-rabbit secondary antibodies (Jackson ImmunoResearch, donkey anti-mouse 715-005-150 and donkey anti-rabbit 715-005-152) in 100 µL of high salt blocking buffer overnight (>16 hours) at 4°C. This was followed by washing with high salt blocking buffer for 5 minutes, three times. Next, the samples underwent embedding and FISH as described above.

After FISH washing, two sets of Epi-PHR hairpins were separately refolded according to the protocol described above. Next, linker oligos for PH1 docking and PH2 docking were hybridized to samples at 40 nM concentration in 100 µL buffer (50% formamide and 20% dextran sulfate in 2xSSC) for 45 min at 37°C and were then washed in 2xSSC with 0.1% v/v Tween-20 in a 60°C water bath for 5 min 3 times. Next, samples were incubated with PH1 hybridization mixture (100 µL refolded first set PH1, 100 µL refolded second set PH1, 200 µL formamide, and 100 µL 50% Dextran Sulfate) for 45 min at 37°C and were then washed in 2xSSC with 0.1% v/v Tween-20 in a 60°C water bath for 5 min 3 times. The samples were next incubated with PH2 hybridization mixture (100 µL refolded first set PH2, 100 µL refolded second set PH2, 80 µL formamide, 120 µL 50% Dextran Sulfate, 20 µL water) for 45 min at 37°C, and were then washed in 35% formamide in 2xSSC with 0.1% v/v Tween-20 for 5 min 3 times. The activators oligos from both sets were hybridized to the sample at 40 nM concentration in 100 µL buffer (25% formamide, 20% Dextran Sulfate, in 2xSSC) for 1 hr at 37°C, and then washed with 35% formamide in 2xSSC with 0.1% v/v Tween-20 for 5 min 3 times. Samples were incubated with H1 hybridization mixture (100 µL refolded first set H1, 100 µL refolded second set H1, 60 µL 5 M NaCl, 140 µL Dextran Sulfate) for 1 hr at 37°C and washed with 35% formamide in 2xSSC with 0.1% v/v Tween-20 for 5 min 3 times. H2 hybridization and washing were performed as for H1. Finally, blocking oligos were hybridized to samples at 100 nM concentration in 400 µL buffer (100 µL HCR buffer, 80 µL 5 M NaCl, 120 µL Dextran Sulfate, and 100 µL water) for 1 hr at 37°C and were then washed with 35% formamide in 2xSSC with 0.1% v/v Tween-20 for 5 min 3 times. The standard digestion protocol was then followed.

Dual-mark Epi-PHR in mouse hippocampus was performed as above, except that concentrations of all PHR components were multiplied by a factor of 2.5X, and the activator step was incubated overnight.

#### Dlk1-Meg3 Epi-PHR

For single-mark Epi-PHR in TSCs, the proximity hybridization reaction and clearing were performed as for cell culture, with the following changes: All washes in a 60°C water bath were replaced with equivalent washes of 55% formamide in 2xSSC with 0.1% v/v Tween-20.

Digestion and clearing were carried out overnight (>16 hr) at 37°C in digestion buffer with 1:100 Proteinase K Solution.

##### Imaging

###### Imaging system

We used a home-built microscope system which has been previously described^104^. In brief, the system consisted of a Nikon Ti2-U body, a Nikon CFI Plan Apo Lambda 60× Oil (NA1.40) objective lens, a 750-nm laser (2RU-VFL-P-500-750-B1R, MPB Communications), a 647-nm laser (2RU-VFL-P-1000-647-B1R, MPB Communications), a 560-nm laser (2RU-VFL-P-1000-560-B1R, MPB Communications), a 488-nm laser (2RU-VFL-P-500-488-B1R, MPB Communications), and a 405-nm laser (OBIS 405 nm LX 50 mW, Coherent). A multi-band dichroic mirror (ZT405/488/561/647/752rpc-UF2, Chroma) on the excitation path directed laser lines to the sample, while the emission path contained a multi-band emission filter (ZET405/488/561/647-656/752 m, Chroma) and a Hamamatsu Orca Flash 4.0 V3 camera. An active auto-focusing system and a motorized x–y sample stage (SCAN IM 112×74, Marzhauser) were used to position samples for imaging. To allow flow of buffers during imaging, samples were loaded into a Bioptechs FCS2 flow chamber. Buffers were directed to the sample through a custom microfluidics system. All microscope systems were centrally controlled from a computer interface using custom software.

###### Color-shift-correction beads

Color-shift-correction beads were prepared fresh for each experiment, as required. For color-shift correction between 560 and 647-nm color channels, we applied TetraSpeck beads (Invitrogen, T7279) at a concentration of 1:200 in Imaging Buffer to a glass coverslip and incubated at RT for 15 min. For color-correction between 647 and 750-nm color channels, biotin-labeled microspheres (Invitrogen, F8767) were applied at a concentration of 1:12000 to a glass coverslip and incubated at RT for 10min, followed by 2x washes in DPBS at RT. The coverslip was then blocked with blocking buffer for 1 hr at RT. The beads were then incubated with streptavidin-dye conjugates (Invitrogen, S32357; Invitrogen, S21384) at a concentration of 1:5000 each in blocking buffer for 15 min at RT with gentle shaking, followed by 3x 3 min washes with DPBS supplemented with 0.1% Tween-20. The beads were finally mounted in imaging buffer, and imaged in a multi-color z-stack, with all correction colors imaged at each z-height before proceeding to the next z-height.

###### Buffers

“Imaging Buffer”^107^ consisted of 50 mM Tris-HCl pH 8.0, 10% wt/v glucose, 2 mM Trolox (Sigma-Aldrich, 238813), 0.5 mg/mL glucose oxidase (Sigma-Aldrich, G2133), and 40 μg/mL catalase (Sigma-Aldrich, C30) in 2xSSC, and was protected from oxidation under a thin layer of mineral oil (Sigma, 330779). “Hybridization Buffer” consisted of 35% formamide (25% for 200 loci experiments) and 0.01% Triton X-100 in 2xSSC. “Wash Buffer” contained 30% formamide (35% for 200 loci experiments) in 2xSSC. “Cleavage Buffer” consisted of 50 mM Tris(2-carboxyethyl)phosphine hydrochloride (Sigma-Aldrich, C4706) and 35% formamide in 2xSSC, adjusted with NaOH (Thermo Scientific, J63736.AK) to pH 7.0. Cleavage buffer was supplemented with blocking oligos corresponding to used readout sequences (sequences are provided in Supplementary Table 1) at 0.5-1.0 μM concentration.

###### Nuclear staining

All samples were incubated with 1:1000 DAPI (Thermo Scientific, 62248) in DPBS prior to imaging. DAPI images were collected in Imaging Buffer using our 405-nm laser immediately at the start of each experiment to maximize signal brightness.

###### Epi-PHR imaging

All images were collected in Imaging Buffer as multi-color z-stacks, with all signal and fiducial colors imaged at each z-height before proceeding to the next z-height. As needed, Epi-PHR signals were then removed by photobleaching in 2xSSC buffer.

###### Sequential FISH imaging

All images were collected as multi-color z-stacks, with all signal and fiducial colors imaged at each z-height before proceeding to the next z-height, to allow for accurate drift correction. At the beginning of each experiment, readouts were hybridized to fiducial markers using protocols equivalent to those described below.

For multiplexed Epi-PHR experiments, each round of readouts was hybridized and cleaved according to the following protocol: Adapter oligos were applied at 50 nM concentration in Hybridization Buffer for 26 min 40 sec and washed in Wash Buffer for 8 min 40 sec. Next, the appropriate dye-labeled readout oligos were applied at 2.5 nM concentration in Hybridization Buffer for 26 min 40 sec and washed in Wash Buffer for 8 min 40 sec. The sample was then rinsed with 2xSSC for 7 min. The sample was then incubated in Imaging Buffer for 7 min before imaging each round. After imaging, the previous round of readout dyes was cleaved and washed away by applying Cleavage Buffer to the sample for 20 min and washing with 2xSSC for 6 min.

For chromatin tracing experiments, the protocol was similar to that described for multiplexed Epi-PHR, except that adapter (30 nM) and readouts (60 nM) were pooled together in Hybridization Buffer and applied to the sample for 27.5 min and washed in Washing Buffer for 4 min 20 sec. The sample was incubated in Imaging Buffer for 4 min before imaging each round. After cleavage with Cleavage Buffer, the sample was washed with 2xSSC for 12 min before hybridizing the readout for the next round of imaging.

For SNP FISH imaging, readout dyes were hybridized to the sample in 20% v/v ethylene carbonate (Sigma-Aldrich, E26258) in 2xSSC for 25 min, and then washed in the same buffer for 3 min 10 sec. The sample was incubated in Imaging Buffer for 3.5 min before imaging. After imaging, SNP FISH signals were removed by photobleaching in 2xSSC buffer.

Readout and adapter oligos were ordered from IDT. Sequences of all readouts and adapters are provided in Supplementary Table 1.

##### Imaging data analysis

All image processing was performed using custom code in MATLAB® software.

###### Color shift correction

Color shift correction was used to account for chromatic aberration between used color channels. This was done by 3D Gaussian fitting the same color-shift-correction beads imaged in two colors and using the MATLAB “cp2tform” function to generate a spatial transformation between these two color channels. This transformation was applied to images using the MATLAB “imtransform” function.

###### Nuclear extraction

Except where noted below, nuclear boundaries used for single-cell and spatial analysis were extracted from DAPI staining images using a customized MATLAB script.

###### Epi-PHR signal-to-background calculations

In the Epi-PHR image, we divided the maximum signal magnitude within a “signal” cuboid by the median signal in a larger “background” cuboid both centered at the same search position. The background cuboid excluded all pixels covered in the x-y plane by the signal cuboid, giving it a “donut” shape. The definitions for the search position, signal cuboid, and background cuboid are given below for each experiment.

In single-molecule FISH (smFISH) experiments, the Epi-PHR search position was defined as 3D position of the smFISH signal (signal: 324x324x600 nm, corresponding to 3x3 pixels in x-y with a pixel size of 108 nm and 3 steps in z with a z step size of 200 nm, background: 0.6 µm in z, and 2x the foci range from gaussian fitting algorithm in x-y). In multiplexed Epi-PHR experiments, the Epi-PHR search position was defined as the median 3D position of bit signals for each decoded gene target molecule (signal: 324x324x600 nm, background: 600 nm in z, and 2x the foci range from gaussian fitting algorithm in x-y). In mouse hippocampus multiplexed Epi-PHR experiments, the Epi-PHR search position was defined as the median 3D position of bit signals (signal: 324x324x600 nm, background: 972x972x600 nm). In chromatin tracing experiments, the Epi-PHR search position was defined as the median 3D position of all tracing loci from which the Epi-PHR signal arose (signal: 324x324x200 nm, background: 2.48x2.48x0.6 µm).

###### Epi-PHR detection efficiency calculation

Detection efficiency in Fig. 2e was calculated as the proportion of signal-to-background ratios above the threshold defined by the 99^th^ percentile of the 2 kb genomic resolution “without primary antibody” control experiment. For Fig. 2g, detection efficiency was similarly calculated using a threshold determined by the 99th percentile of signal-to-background ratios from the IgG control experiment.

###### Decoding of 200-gene single-dual-mark Epi-PHR

To detect multiplexed FISH signals in cell nuclei, we began by segmenting the nuclei using DAPI staining and excluded overlapping nuclei based on the circularity of the DAPI contour. FISH signals within the segmented nuclei were fitted to 3D Gaussian functions to determine their center positions along the x, y, and z axes. To correct for sample drift between hybridization rounds, we applied the same algorithm to the genomic fiducial mark images, determined sample movement from the fiducial mark positions, and subtracted these movements from the raw foci positions. Subsequently, all identified FISH signals were pooled, and pairwise spatial distances were measured between individual FISH signals. If the distance between FISH signals was smaller than 2 pixels (216 nm) in the x and y directions and 2 z-steps (400nm) in the z direction, the bit information was extracted and used to construct the barcode. The constructed barcode was decoded into the targeted genomic locus using a 36-bit HD2 code, with only perfectly matching barcodes being assigned to specific genomic loci.

###### Decoding mouse hippocampus Epi-PHR

Decoding of Epi-PHR data from the mouse hippocampus followed the same analytical pipeline as described above, with modifications to the nuclei segmentation and FISH foci detection steps. Nuclei were segmented based on DAPI staining using Cellpose 3.0^108^ with the ’nuclei’ model, employing the parameters: stitch_threshold = 0.4, flow_threshold = 0.7, and cellprob_threshold = -9. FISH signals within the segmented nuclear volumes were subsequently identified and localized using a three-dimensional radial center algorithm to determine their centroid coordinates along the x, y, and z axes^109^.

###### Epi-PHR signal rate calculations

For each profiled epigenetic mark, all measured Epi-PHR signal-to-background values were pooled and fitted to a Gaussian mixture distribution model with two components using the MATLAB "fitgmdist" function. The mean value of the second component served as the signal-to-background threshold for Epi-PHR signal rate calculations. To determine the Epi-PHR signal rate for each targeted locus, the number of Epi-PHR measurements above the threshold was counted and divided by the total number of decoded FISH foci for that locus.

###### Hippocampus Epi-PHR enrichment Z-score calculations

Anatomical brain regions were manually segmented by aligning DAPI-stained images with corresponding reference sections from the Allen Brain Atlas (http://developingmouse.brain-map.org/static/atlas). For each segmented region, the mean Epi-PHR signal-to-background ratio was computed for each target gene. Subsequently, Z-scores were calculated independently for each histone modification mark. The final enrichment score for each gene locus was determined by summing the Z-scores from the two histone marks. The genes in the heatmap shown in Fig. 4m were ordered using the following procedure: (1) gene loci exhibiting the largest Z-score differences across the defined regions were grouped; (2) within each group (CA-enriched and DG-enriched), the top 20 loci with the greatest differential enrichment were ranked; and (3) these groups were combined and arranged according to the anatomical regions presented in the figure.

###### Chromatin tracing

As a first step in processing chromatin tracing datasets, we corrected the spatial drift among all rounds of imaging by 3D Gaussian fitting fiducial marker foci in each multi-color image stack.

Next, we used a 3D Gaussian function to fit chromatin tracing foci in all images. Chromatin traces were assembled from fit tracing foci by sequentially adding foci to nearby previously identified traces, assigning unmatched foci as new traces. Traces containing more than a threshold number of loci were retained for downstream analyses.

###### Dlk1-Meg3 locus Epi-PHR and chromatin tracing analysis

To identify the parent-of-origin of each assembled trace, we first removed background signal from drift- and color-corrected SNP FISH images by subtracting a copy of the image stack processed by image opening. The parent-of-origin-specific SNP FISH signal of each trace was calculated as the maximum signal magnitude in a 1.40x1.40x1.40 µm cuboid centered on the median 3D position of that trace. For each image, SNP FISH signals were then normalized to the highest calculated SNP FISH signal in that image. To assign a parent-of-origin to a trace, that trace was required to have SNP FISH signals of >= 0.1 for the assigned parent-of-origin, and <= 0.07 for the opposite parent-of-origin.

Boundary scores at the edge of tracing loci in median spatial distance matrices were calculated as shown in Extended Data Fig. 8a. Mirrored above and below the diagonal, a sliding window of pixels extending 5 pixels was divided into the neighboring group of pixels. Boundary score was calculated as the mean inverse of this division. Maternal contact bias for each tracing locus in median spatial distance matrices were calculated as shown in Extended Data Fig. 8b. All contacts for each locus in the maternal matrix were divided by the same contacts in the paternal matrix.

Maternal contact bias was calculated as the mean inverse of this division along each column. Dimensional reduction by *t*-Distributed Stochastic Neighbor Embedding (t-SNE) ^91^ normalized by cosine distance was performed on traces based on the pairwise spatial distances among their tracing loci. Traces were only included if they had at least 80% of tracing loci detected and were not missing more than 2 consecutive tracing loci. Missing pairwise spatial distance values were filled in by linear interpolation for t-SNE.

For single-cell and spatial analyses, nuclear boundaries were extracted manually using as reference a max projection of DAPI signal magnitude gradient, referring to no other images or data to avoid potential bias. Cells with substantial overlap (greater than ∼15%) were rejected. Distance to the nuclear boundary for each trace was calculated as the distance in the x-y plane to the nearest nuclear boundary pixel. *Meg3* loci were considered to be in contact with the nuclear boundary if their nearest boundary distance was <= 200 nm.

##### Epigenomic MERFISH

Epigenomic MERFISH was performed according to the published protocol^44^ in IMR90 cells, with the addition of the RNase treatment step described above to remove endogenous RNA.

###### Epigenomic MERFISH 200-loci profiling

Epigenomic MERFISH was performed using the 200-loci library. Imaging was performed as above, with the addition of 0.05% murine RNase inhibitor (New England BioLabs, M0314) in all imaging buffers to prevent degradation of *in vitro* RNA transcripts.

###### Epigenomic MERFISH signal rate calculations

To determine the epigenomic MERFISH signal rate for each targeted locus, the total number of decoded epigenomic MERFISH foci were counted and divided by the total number of cells.

###### Epigenomic MERFISH and Chromatin Tracing

Epigenomic MERFISH was performed using the library described above for Epi-PHR with chromatin tracing in cell culture. During imaging, buffers were supplemented with 0.05% murine RNase inhibitor (New England BioLabs, M0314) to prevent degradation of *in vitro* RNA transcripts. The PH1 docking site on all probes was used to generate epigenomic MERFISH signals. After epigenomic MERFISH data was collected, the sample was removed from the microscope for the chromatin tracing FISH protocol. The sample was digested with RNase and then submerged in pre-hybridization buffer, as described above. The sample was then re-hybridized to the Epi-PHR with chromatin tracing in cell culture library according to the following protocol: the sample was denatured in a water bath at 84°C for 3 min and incubated at least 16 hr at 37°C. The sample was then washed at RT 2 times for 15 min each with 55% formamide in 2xSSC supplemented with 0.1% v/v Tween-20. The sample was reloaded onto the microscope and realigned to the previously selected FOVs using the DAPI staining pattern.

Sequential FISH imaging was then performed as above. In this experiment, fiducial markers were hybridized to the chromatin tracing library primer binding sites.

##### Sequencing

###### ChIP-Seq Analysis

Processed IMR90 ChIP-Seq data for H3K36me3, H3K27me3, H3K9me3, H3K9ac, DNase, and H3K27ac were downloaded from the Roadmap Epigenomics Project (https://egg2.wustl.edu/roadmap/data/byFileType/peaks/consolidated/narrowPeak/, accession E017-H3K36me3, E017-H3K27me3, E017-H3K9me3, E017-H3K9ac, E017-Dnase and E017-H3K27ac) as narrow peaks^48^. For each targeted locus and epigenetic mark, ChIP-seq peak density was calculated by summing the narrow peak heights over the targeted region and normalizing by the genomic length of that region. For the categorized Epi-PHR signal rates analysis (Figs. 3h, 3i, 4g, and 4h), loci with a ChIP-seq density greater than zero were classified as high ChIP-seq density loci. The remaining loci were grouped into low ChIP-seq density loci.

###### RNA-Seq Library Preparation

TSCs were collected and frozen at -80°C until use. RNA was extracted with RNeasy micro kit’s (Qiagen, 74004) standard cell culture RNA protocol with DNase digestion. The quality of RNA samples was measured using the Agilent TapeStation RNA ScreenTape and Reagents (5067-5579); all samples had RIN > 8.8. RNA libraries were generated using the Takara SMARTer® Stranded Total RNA-seq Kit v3-Pico Input Mammalian and the 24U dual index kit per manufacturer specifications using the direct total RNA input protocol (Takara Biosciences, 634485). Library size was confirmed via the Agilent TapeStation DNA ScreenTape to meet Takara’s guidelines for sequencing (5067-5582). Sequencing was performed by the Yale Center for Genomic Analysis and resulted in between 232M and 310M reads. Sequencing was performed with 150 bp paired-end reads on an Illumina NovaSeq 6000.

###### SNP Identification

To phase RNA-seq data and generate SNP sensitive FISH probes for C57/CAST F1 hybrid cells, we generated a list of high-quality SNPs that are homozygous within C57 or CAST and yet distinct between the C57 and CAST genomes. In brief, we used the mouse genome project (mgp.v5.merged.snps_all.dbsnp142.vcf), and utilized custom Python scripts to identify high quality (MGP quality flag = “PASS”) homozygous SNPs between the CAST genome and the C57 genome (Reference + C57 homozygous SNPs), and output those to a custom filtered VCF file. Using this framework, we identified 17,491,436 SNPs homozygous in both the C57 and CAST lineages that passed quality control checks (>99% probability of SNP) that we could use for phasing reads. We used GATK FastaAlternateRefereranceMaker with “--snp-mask” on the GRCm38.fa genome against these sites to generate an N masked mm10 genome to prevent reference sequence alignment bias to the C57 line^110^. Individual Fasta Genomes for C57 and CAST were also generated using GATK FastaAlternateReferanceMaker from this VCF file for downstream FISH probe generation.

###### Phased RNA-Seq Analysis

Raw reads underwent QC analysis by FASTQC (http://www.bioinformatics.babraham.ac.uk/projects/fastqc/). Demultiplexed FASTQs were had their UMI’s extracted using UMI_Tools (discarding the 6 bp spacer per Takara’s manual), and reads were aligned to an “N-Masked” GRCm38 genome using STAR^111^ using “--twopassMode Basic, --alignEndsType EndToEnd, --outSAMattributes NH HI NM MD”. Aligned reads were then UMI deduplicated using UMI_Tools, where per the Takara SMARTer® Stranded Total RNA-seq Kit v3-Pico Input Mammalian kit instructions, each UMI specifies a single strand of template RNA, with the flags “—method=unique, --unpaired-reads=discard, --paired”^112^. BAM files were sorted as needed using BAMtools v2.5.1^113^ and read counts at exons were quantified using HTSeq-count v0.9.1 “—additional-attr=genename, -r pos, -t exon, -I gene_id, --stranded=reverse” using the GRCm38.102.gtf file^114^. Data preparation was performed in Python. Statistics and figure preparation were performed in R (v4.2.2) and Python.

Bam files were split by allele using SNPSplit v0.3.4^115^. In brief, SNPSplit uses a VCF file (generated above) and the original N-mask-aligned BAM file and splits it into three output files, one for the paternal genome, one for the maternal genome, and one for unidentified reads. For a read to be called as paternal or maternal, a SNP must lie in the sequenced read area (150 bp per read, 300 bp/read pair). A single SNP in one paired-end read is sufficient to split both reads.

Allelic contributions were calculated in R (v4.2.2) using DESeq2 (v1.38.3) with FDR pValueAdjustment^116^, with a minimum count per gene of 30 for downstream analysis. Imaging of pileups was performed in IGV without windowing and presented with liftover coordinates for mm39.

###### CUT&RUN Library Preparation

Fresh MEF Depleted TSCs were quantified and split into 200,000 cells per condition for downstream CUT&RUN reactions. CUT&RUN was performed using the EpiCypher CUTANA CUT&RUN kit (14-1048, v3.1) per the manufacturer’s specifications with the following antibodies: IgG Control (CUTANA Kit Rabbit IgG Negative Control Antibody), H3K9me3 (abcam, ab8898), and H3K4me3 (CUTANA Kit, Positive Control). DNA was then converted to a sequencing library using the CUT&RUN Library Prep Kit (14-1001, v1.0) according to the manufacturer’s specifications. Libraries were run on Agilent TapeStation DNA ScreenTape to check for library size specificity of 200-700 bp (5067-5582). Sequencing was performed with 150 bp paired-end reads on an Illumina NovaSeq 6000.

###### CUT&RUN Analysis

Sequencing resulted in ∼50M reads per sample, a higher number than usually used in CUT&RUN analysis due to the lossy nature of SNP-splitting in phased analysis. Reads were analyzed via FastQC (http://www.bioinformatics.babraham.ac.uk/projects/fastqc/) and trimmed via Trimmomatic^117^. Trimmed reads were aligned to an N-masked aligned C57/Cast genome (mm10) using Bowtie2 with “--local -N 1” and paired-end input^118^. Aligned reads were converted to BAM, sorted and deduplicated using SAMtools v1.11^113^. SNPsplit was used to isolate paternal and maternal reads with the “—paired” flag. Reads that could not be assigned to a parent of origin were discarded for downstream analysis. ∼8% (7.76%-10.76%) of reads were assignable to either the maternal or paternal genome for downstream processing. Peaks were called on split bam files using MACS2 with “-g mm -f BAMPE –nomodel –shift -160 –extsize 150 --broad” settings for H3K9me3, with IgG as an input control. Samples were run with their phased input control (i.e. Replicate1_Maternal & IgG_Maternal, Replicate1_Paternal & IgG_Paternal)^119^. BigWig signals for visualization were obtained via deeptools bamCoverage^120^ normalized to read depth (--binSize 10 --normalizeUsing CPM --extendReads 300 –centerReads). Imaging of pileups was performed in IGV without windowing and presented with liftover coordinates for mm39^121^.

## Statistical Analysis

Sample sizes are available in figure legends. All relevant statistical tests were two-sided.

## Data availability

Sequencing data are available via Gene Expression Omnibus accession GSE274718 (token: ipwfugisfjijfyv). Raw imaging data are available from the corresponding author upon request and are not deposited online due to prohibitive size. Processed imaging datasets are available at https://campuspress.yale.edu/wanglab/EpiPHR.

## Code availability

Original code used within this study are available at https://campuspress.yale.edu/wanglab/EpiPHR. Open-source codes for imaging data collection are available at https://github.com/ZhuangLab/storm-control/.

## Acknowledgements

We thank Dr. Mauro Calabrese for the gift of the trophoblast stem cell line (C/B, 6-2). We thank all members from Wang and Xiao labs for helpful discussions. This work was supported by the National Human Genome Research Institute (R01HG011245), the National Institutes of Health Cellular Senescence Program (UG3/UH3CA268202), the National Institute of Child Health and Human Development (R01HD102533), the National Institute of General Medical Sciences (R35GM136346), and the Carol and Eugene Ludwig Family Foundation. J.S.D.R. was supported by NIH Predoctoral Training Grant (2T32GM007499). M.H.A. was funded by the National Science Foundation Graduate Research Fellowship (DGE1752134), the Gruber Foundation, and the KS & Feili Lo Foundation. Sequencing research reported in this publication was in part supported by the National Institute of General Medical Sciences of the National Institutes of Health under Award Number 1S10OD030363-01A1.

## Author contributions

S.W. conceived the study; Y.Chen, J.S.D.R. and S.W. designed and tested the Epi-PHR system; Y.Chen, J.S.D.R., and M.H.A. performed experiments and analyzed data, with the help of S.J., Y.Cheng, Y.Z. and M.L.; Y.Chen, J.S.D.R., and M.H.A. wrote the first manuscript; S.W. and A.X. supervised the study; all authors revised and approved the final manuscript.

## Ethics declarations

### Supplementary information

Supplementary Table 1. Oligo sequences of Epi-PHR components, readout adapters and probes, blocking oligos, primers, and FISH probe template libraries.

Supplementary Table 2. Codebook for 200-gene Epi-PHR experiment.

Supplementary Table 3. Gene list and codebook for 200-gene Epi-PHR in mouse hippocampus.

Supplementary Table 4. Data values displayed in figures.

---

**Extended Data Fig. 1.**
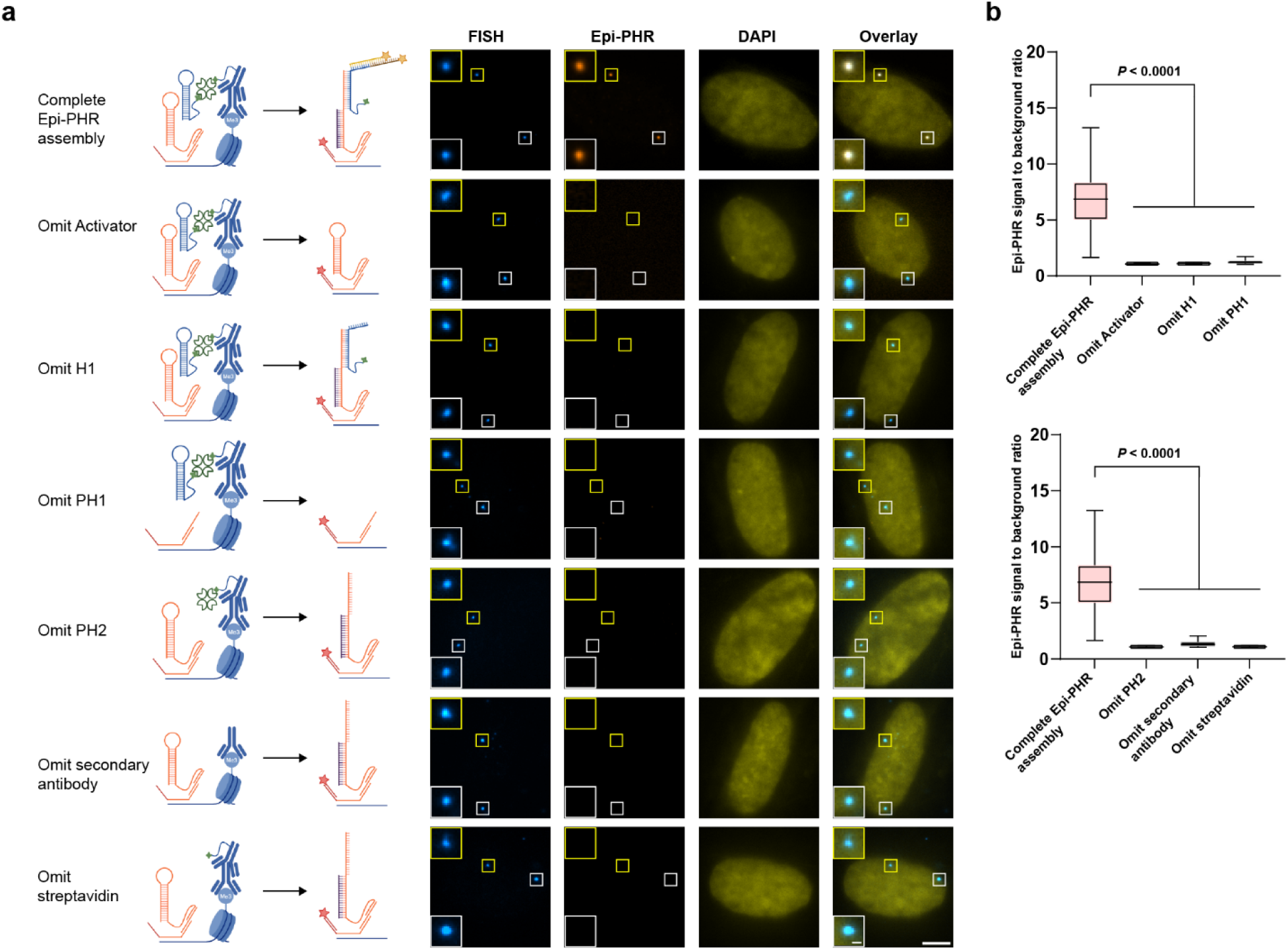
| Performance assessment of the Epi-PHR hairpins. **a**, (Left) Illustrations depicting the experimental design and the anticipated H3K9me3 Epi-PHR assembly following digestion. (Right) Representative 100-kb H3K9me3 FISH, Epi-PHR, and DAPI images from different conditions. FISH, Epi-PHR, and DAPI were imaged from different color channels and merged into one image. The corner panels show magnified images that use FISH signals as center positions. Scale bar for the original images: 5 µm; scale bar for the magnified images: 500 nm. **b**, Signal-to-background ratios of Epi-PHR measured at FISH positions. In box plots, central lines represent median values, boxes show interquartile ranges (IQRs), and whiskers indicate non-outlier minimum and maximum values. Outliers were defined as values further than 1.5 times the IQR away from the box. Statistical significance was determined by unpaired two-tailed Mann-Whitney test. Sample sizes are as follows: complete Epi-PHR assembly (n = 262), omit activator (n = 231), omit H1 (n = 253), omit PH1 (n = 212), omit PH2 (n = 215), omit secondary antibody (n = 252), omit streptavidin (n = 216).

**Extended Data Fig. 2.**
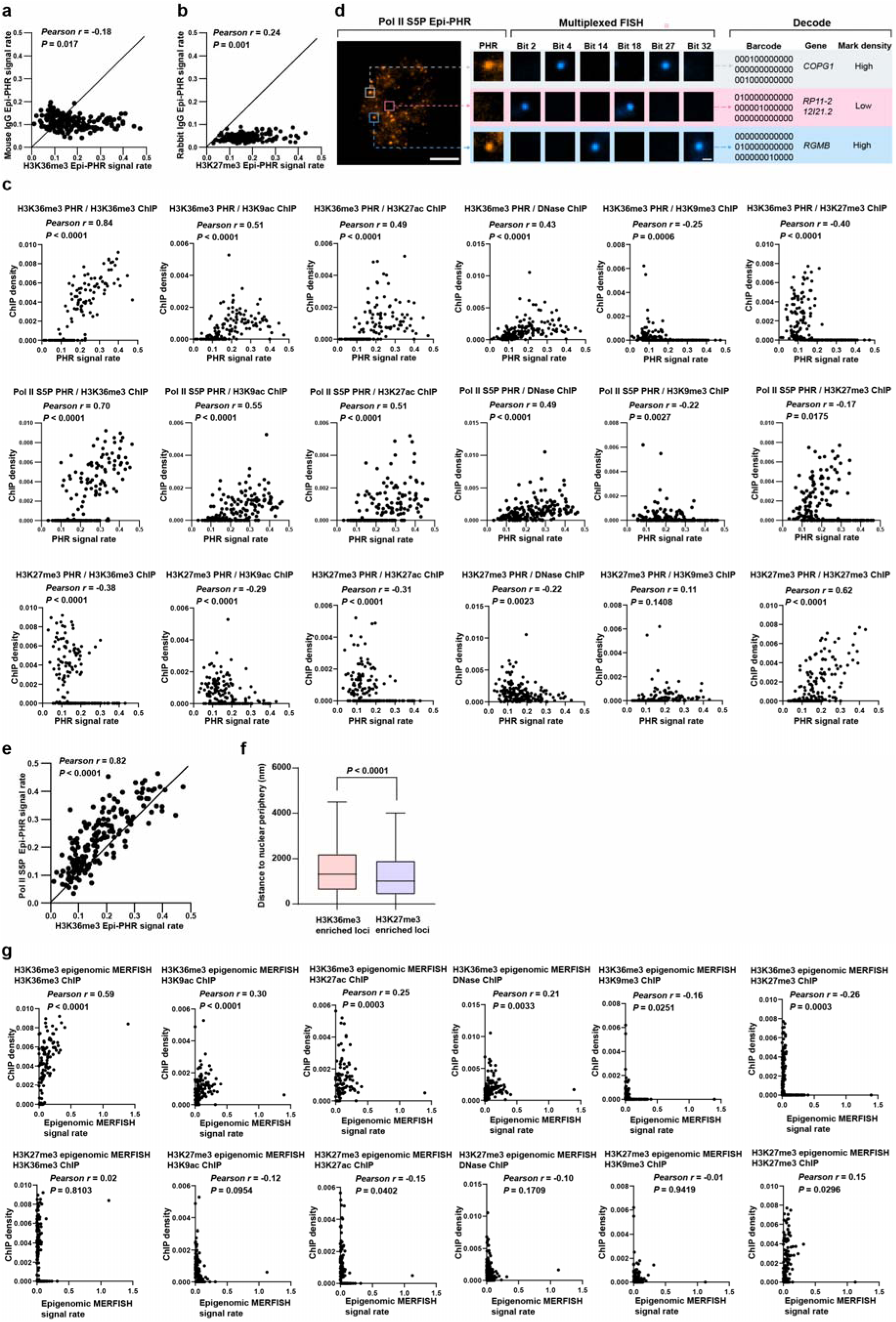
**| Additional analyses from 200-gene Epi-PHR profiling experiments. a-b**, Epi-PHR signal rate correlations between H3K36me3 and mouse IgG, and between H3K27me3 and rabbit IgG. Pearson correlation coefficients and statistical significance determined by two-tailed tests are shown. **c**, Detailed correlation plots for comparing H3K36me3, Pol II S5P, and H3K27me3 Epi-PHR signal rates with H3K36me3, H3K9ac, H3K27ac, DNase, H3K9me3, and H3K27me3 ChIP-seq densities^48^. **d**, Representative 200-gene Pol II S5P Epi-PHR and multiplexed FISH images. Magnified images display three decoded gene loci with their associated Epi-PHR signals and multiplexed FISH signals in selected rounds of readout hybridization. Scale bar for the original images: 5 µm; scale bar for the magnified images: 500 nm. **e**, Epi-PHR signal rate correlation between H3K36me3 and Pol II S5P profiling experiments. Pearson correlation coefficient and statistical significance determined by a two-tailed test are shown. **f**, Distances between H3K36me3 or H3K27me3 enriched loci and nuclear peripheries, defined by DAPI signal boundaries. For each cell, Z-scores were calculated based on Epi-PHR signal-to-background ratio, and loci with Z-scores greater than 2.5 were selected for distance measurements. In box plots, central lines represent median values, boxes show interquartile ranges (IQRs), and whiskers indicate non-outlier minimum and maximum values. Outliers were defined as values further than 1.5 times the IQR away from the box. Statistical significance was determined by an unpaired two-tailed Mann-Whitney test. Sample sizes are as follows: H3K36me3 enriched loci (n = 2946), H3K27me3 enriched loci (n = 3645). **g**, Detailed correlation plots for comparing H3K36me3 and H3K27me3 epigenomic MERFISH signal rates with H3K36me3, H3K9ac, H3K27ac, DNase, H3K9me3, and H3K27me3 ChIP-seq densities^48^.

**Extended Data Fig. 3.**
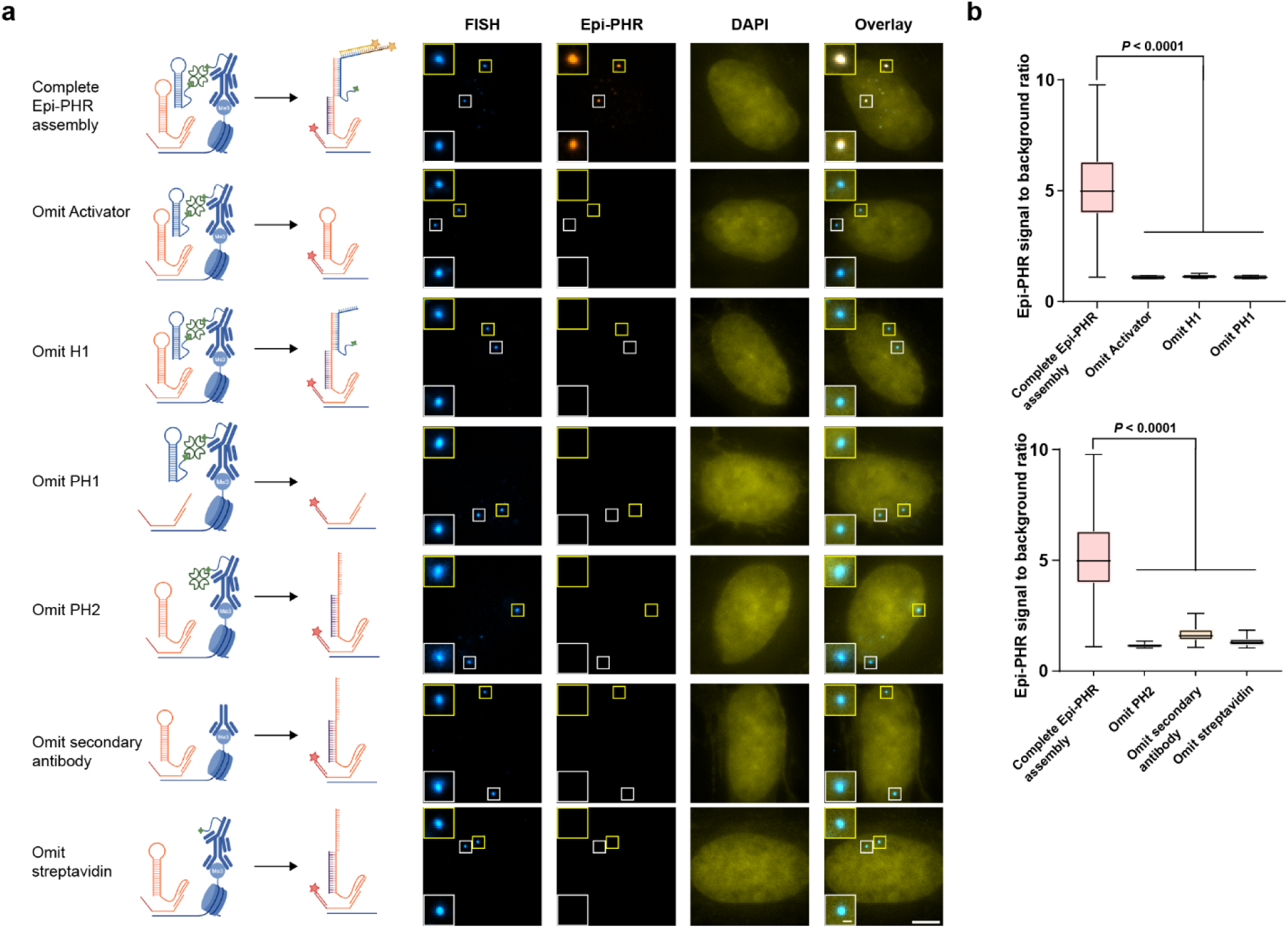
| Performance assessment of the second set of Epi-PHR hairpins. **a**, (Left) Illustrations depict the experimental design and the anticipated H3K9me3 Epi-PHR assembly following digestion. (Right) Representative 100-kb H3K9em3 FISH, Epi-PHR, and DAPI images from different conditions. Corner panels show magnified images that use FISH signals as center positions. Scale bar for the original images: 5 µm; scale bar for the magnified images: 500 nm. **b**, Signal-to-background ratios of Epi-PHR measured at FISH positions. In box plots, central lines represent median values, boxes show interquartile ranges (IQRs), and whiskers indicate non-outlier minimum and maximum values. Outliers were defined as values further than 1.5 times the IQR away from the box. Statistical significance was determined by unpaired two-tailed Mann-Whitney test. Sample sizes are as follows: complete Epi-PHR assembly (n = 426), omit activator (n = 320), omit H1 (n = 278), omit PH1 (n = 295), omit PH2 (n = 251), omit secondary antibody (n = 337), omit streptavidin (n = 255).

**Extended Data Fig. 4.**
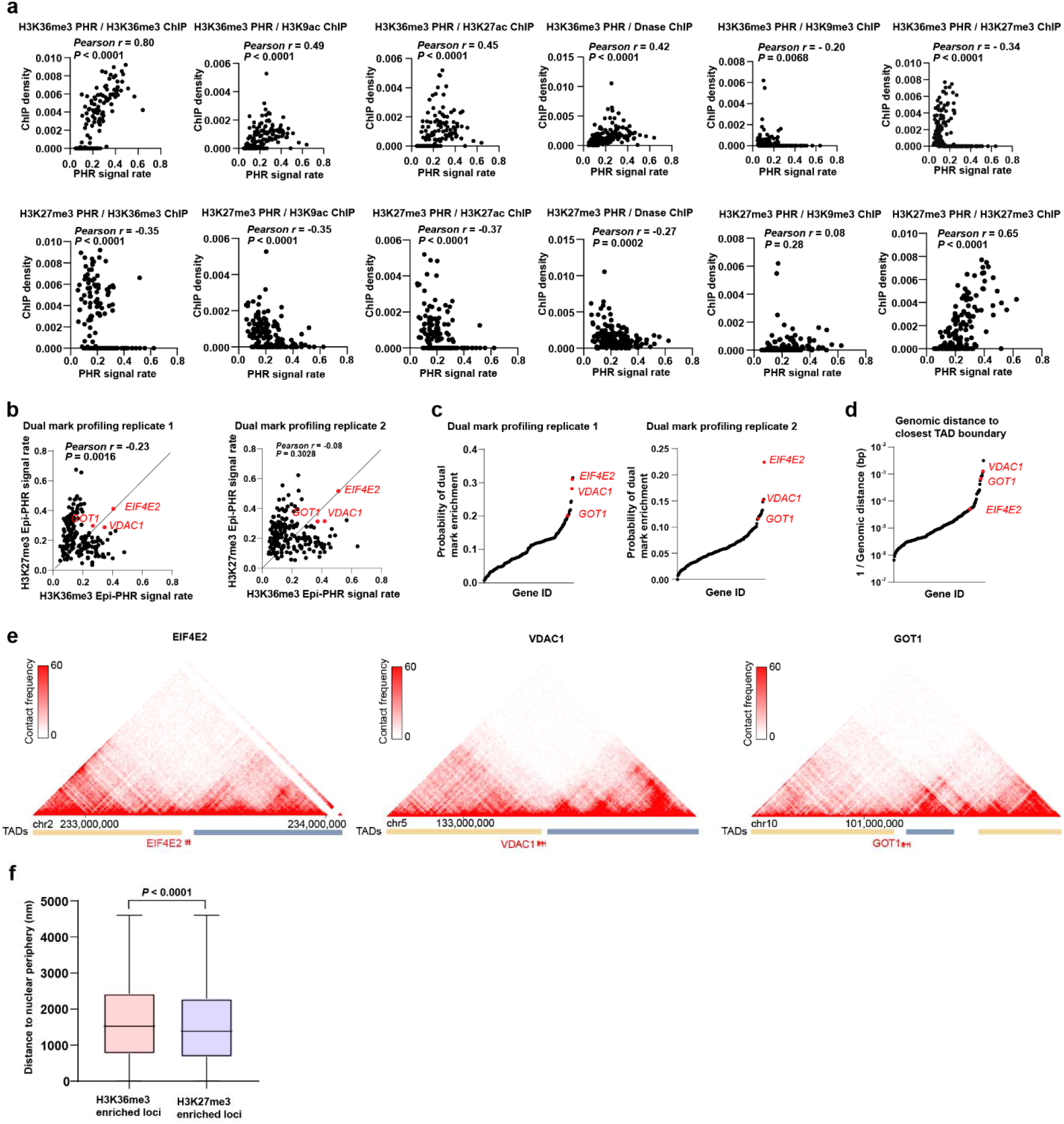
| Additional analyses from 200-gene dual-mark Epi-PHR profiling experiments. **a**, Detailed correlation plots for comparing dual-mark H3K36me3-H3K27me3 Epi-PHR signal rates with H3K36me3, H3K9ac, H3K27ac, DNase, H3K9me3, and H3K27me3 ChIP-seq densities^48^. **b**, Epi-PHR signal rate correlation between H3K36me3 and H3K27me3 in dual-mark profiling experiments. Pearson correlation coefficient and statistical significance determined by a two-tailed test are shown. **c**, Probability of dual-mark enrichment for each targeted locus. **d**, Genomic distances of targeted genes to closest TAD boundaries. **e**, Hi-C contact frequency maps^21,122^ near select genes with high H3K27me3 and H3K36me3 dual-mark enrichment probabilities. **f**, Distances between H3K36me3 or H3K27me3 enriched loci and nuclear peripheries, defined by DAPI signal boundaries. For each cell, two sets of Z-scores were calculated based on H3K36me3 or H3K27me3 Epi-PHR signal-to-background ratios. Loci were selected for distance measurements if they had Z-scores greater than 2.5 in one channel and less than 0 in the other channel. In box plots, central lines represent median values, boxes show interquartile ranges (IQRs), and whiskers indicate non-outlier minimum and maximum values. Outliers were defined as values further than 1.5 times the IQR away from the box. Statistical significance was determined by an unpaired two-tailed Mann-Whitney test. Sample sizes are as follows: H3K36me3 enriched loci (n = 1369), H3K27me3 enriched loci (n = 1331).

**Extended Data Fig. 5.**
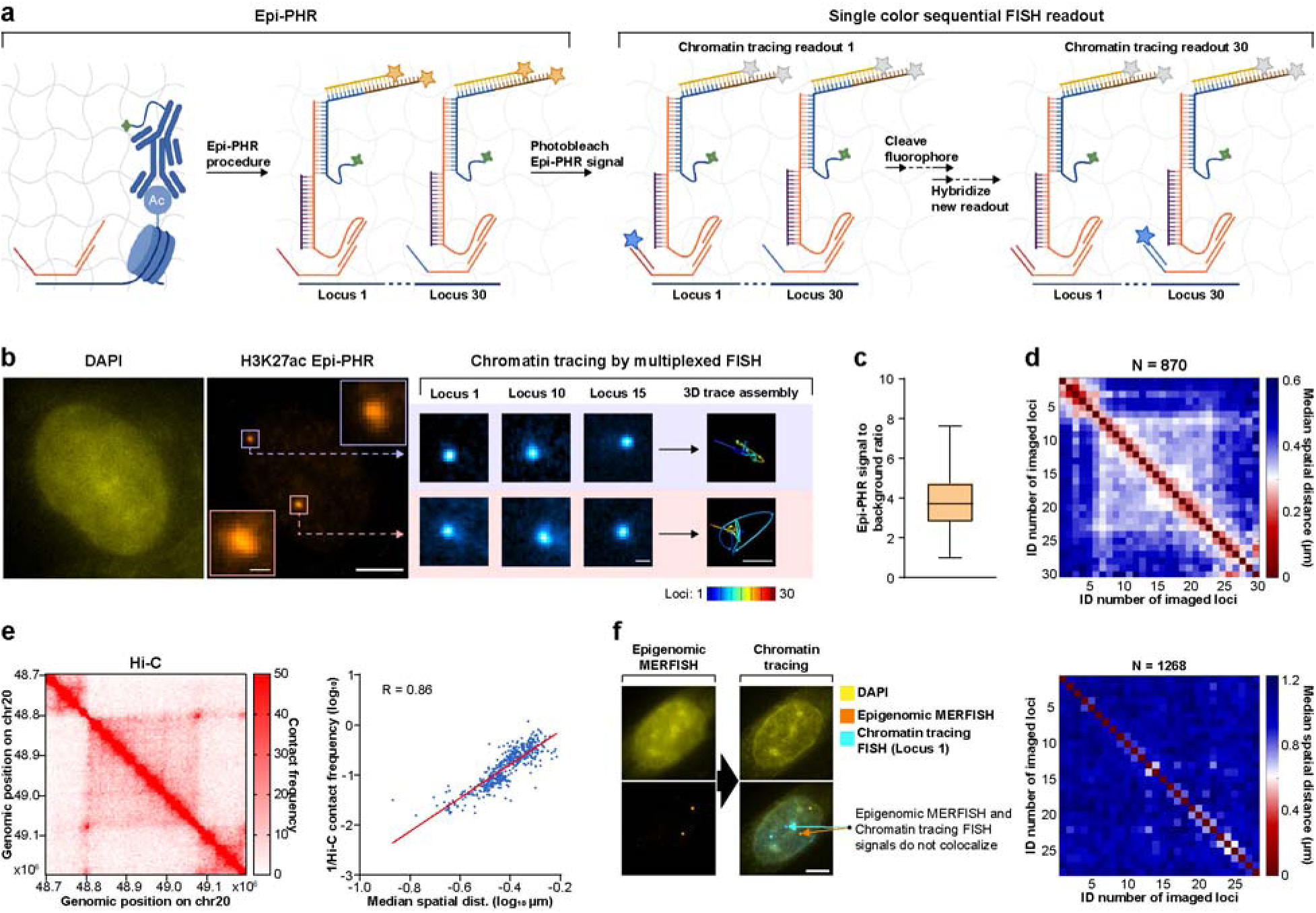
| Epi-PHR combined with chromatin tracing. **a**, Experimental design of Epi-PHR combined with chromatin tracing. PH1 hybridizes to all FISH probes via an adapter. During imaging, Epi-PHR signals were recorded and then photobleached. The spatial positions of tracing loci were then sequentially visualized through 30 rounds of single-color imaging. **b**, Representative H3K27ac Epi-PHR and chromatin tracing images. Sequential FISH foci are assembled into 3D chromatin traces. Scale bars for original images: 5 µm; scale bars for magnified images: 500 nm. **c**, Strong Epi-PHR signals colocalize with 3D chromatin traces. The central line represents the median value, the box shows the interquartile range (IQR), and whiskers indicate non-outlier minimum and maximum values. Outliers were defined as values further than 1.5 times the IQR away from the box. **d**, Median inter-loci spatial distances from chromatin tracing, where each matrix element represents the median distance between two tracing loci. Tracing spanned 500 kb in 30 loci (∼16.6 kb resolution). **e**, (Left) Reanalyzed Hi-C contact frequencies across the chromatin tracing target region (5 kb resolution^21^). (Right) Correlation between inverse Hi-C contact frequency and chromatin tracing median spatial distances. Hi-C data were calculated by taking the mean of all 5-kb resolution Hi-C contact frequencies entirely within the coordinates of each pair of tracing loci. **f**, Epigenomic MERFISH^44^ and chromatin tracing of the same 500-kb region as in D. (Left) Epigenomic MERFISH signals do not colocalize with DNA FISH signals. (Right) Chromatin tracing after epigenomic MERFISH retains no expected chromatin folding. (NOTE: Tracing loci 16 and 30 shown in D were not imaged in F, so the total number of tracing loci was 28.)

**Extended Data Fig. 6.**
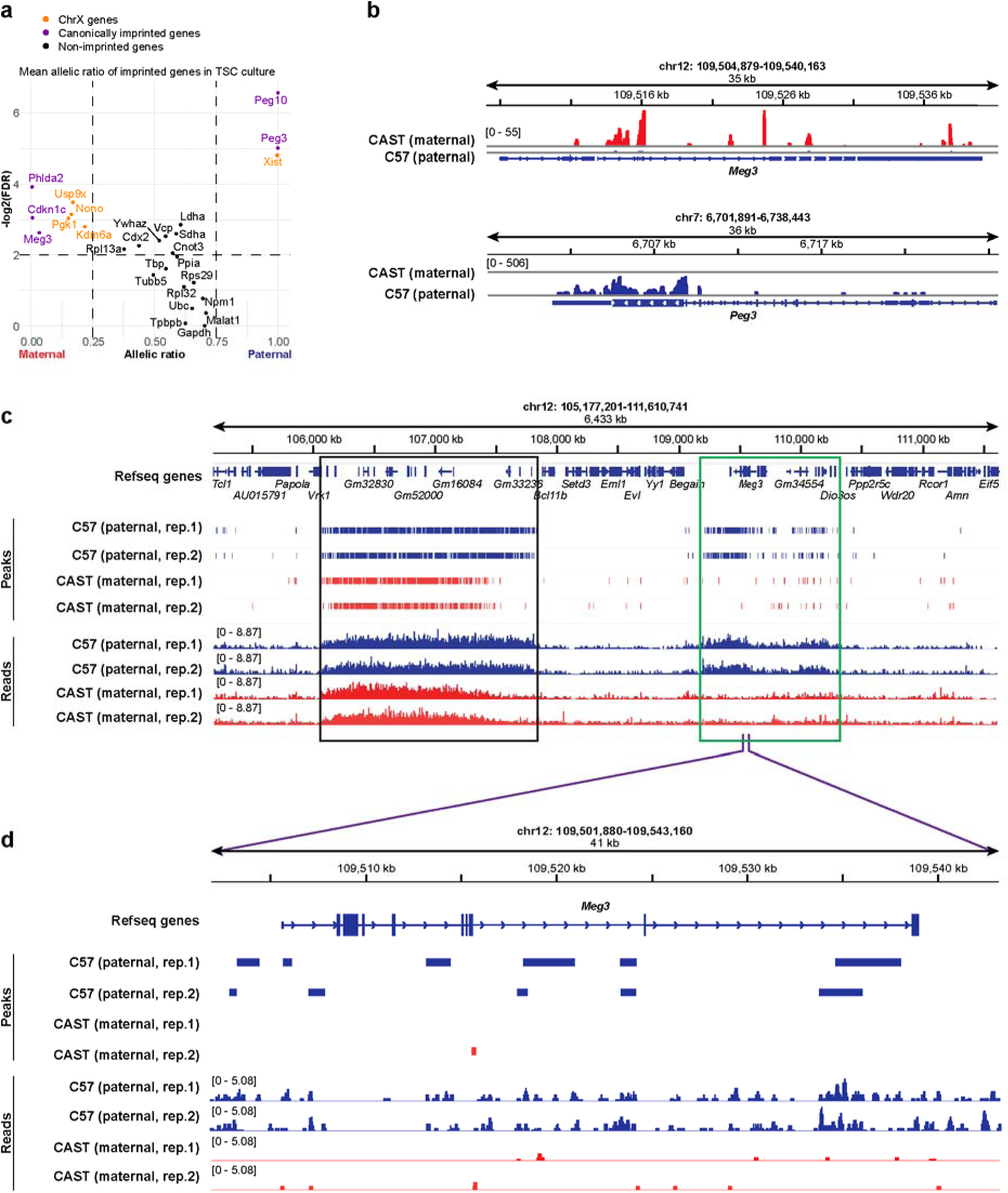
| Phased RNA-seq and CUT&RUN of trophoblast stem cells (TSCs). **a**, Allelic frequency of known imprinted genes (purple), non-imprinted genes (black), and X-Chromosome genes (orange) in TSCs. Note that in the murine placenta, X-chromosome expression is uniformly maternal, with the exception of the noncoding RNA *Xist* which is paternally expressed (see Ref. ^123^). Vertical and horizonal cutoffs are at allelic ratio = 0.25 and 0.75, and -log2(FDR) = 2, respectively. **b**, Phased RNA-seq reads of (top) *Meg3*, showing maternal expression bias, and (bottom) *Peg3*, showing paternal expression bias. **c**, Phased H3K9me3 CUT&RUN in TSCs. At the Mb scale, H3K9me3 can be seen as biallelically deposited at non-imprinted loci (black box) but enriched on the paternal chromosomes at the *Meg3* Imprinted locus (green box). **d**, Phased H3K9me3 peaks and reads are enriched on the paternal allele of the maternally expressed *Meg3* locus.

**Extended Data Fig. 7.**
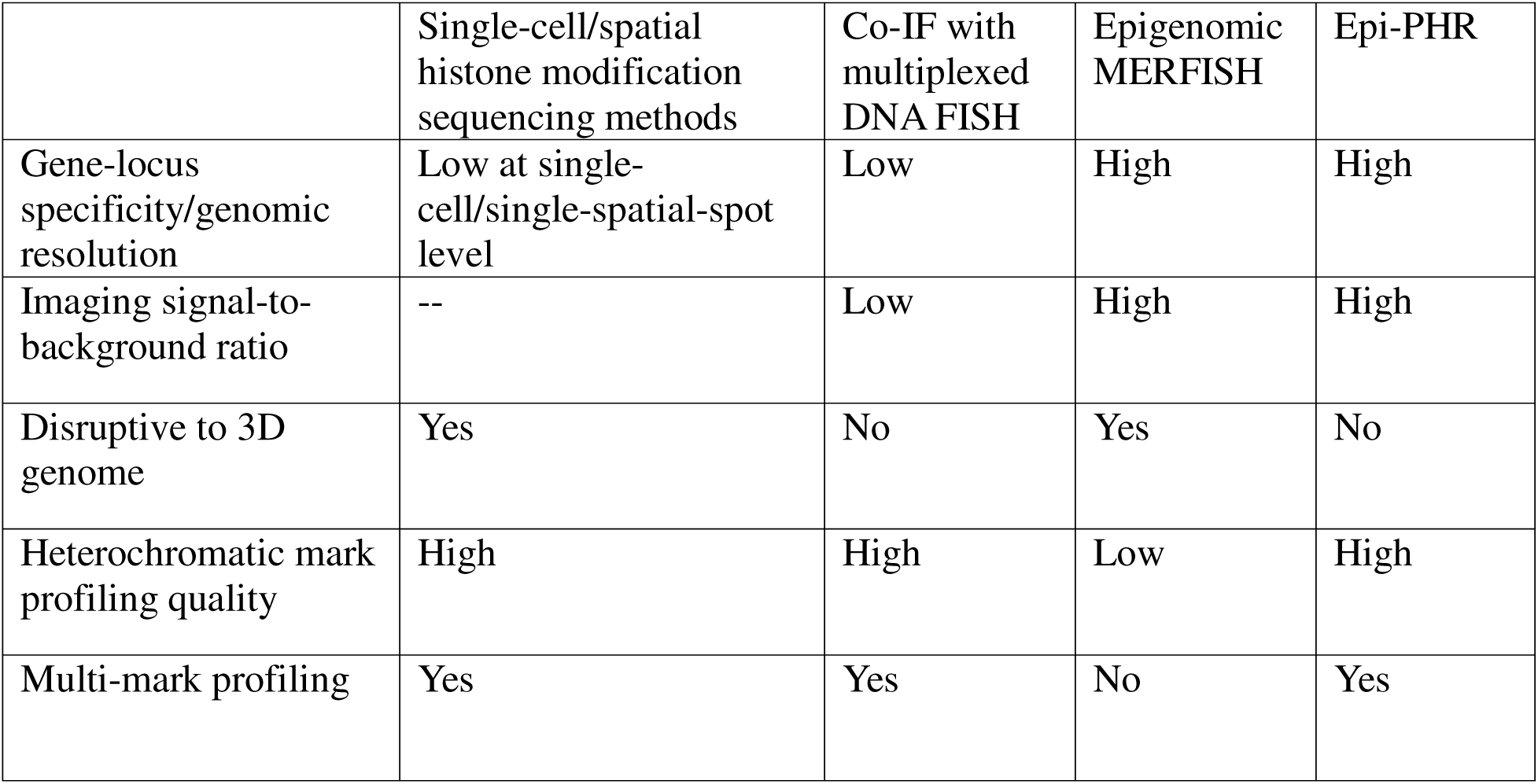
| Comparison of Epi-PHR to other single-cell/spatial epigenetic mark profiling methods.

**Extended Data Fig. 8.**
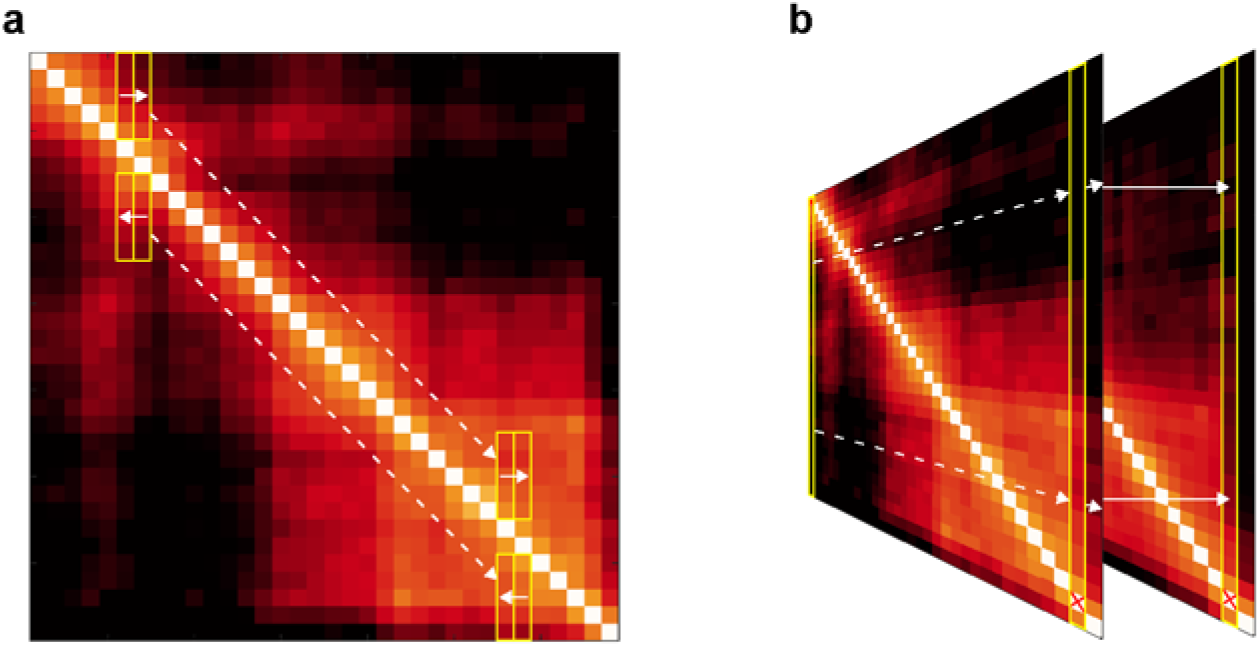
| Methods for calculating chromatin domain boundary score and maternal chromatin contact bias. **a**, To calculate the boundary score at the edge of each tracing locus, a sliding window of pixels (yellow rectangles, dashed white arrows) above and below the diagonal was divided by a neighboring group of pixels (solid white arrows), and the inverse mean of the result was taken. **b**, To calculate the maternal contact bias, each column in the median spatial distance matrix (yellow rectangles, dashed white arrows), with the on-diagonal pixel removed, was extracted from the maternal and paternal matrices and used in a division (maternal column divided by the corresponding paternal column [solid white arrows]). The inverse mean of the result was taken as the bias of each chromatin tracing locus.

